# miniMTI: minimal multiplex tissue imaging enhances biomarker expression prediction from histology

**DOI:** 10.64898/2026.01.21.700911

**Authors:** Zachary Sims, Sandhya Govindarajan, Kaoutar Ait-Ahmad, Cigdem Ak, Marigold Kuykendall, Gordon B. Mills, Ece Eksi, Young Hwan Chang

**Author notes:** co-corresponding Contact: Young Hwan Chang. These authors contributed equally to this work (Zachary Sims, Sandhya Govindarajan, Kaoutar Ait-Ahmad).

## Abstract

Virtual multiplexing from routine histology has advanced rapidly, yet morphology alone provides limited access to molecular state, imposing an intrinsic ceiling on H&E-only inference. Here, we introduce miniMTI, a molecularly anchored virtual staining framework that determines the minimal set of experimentally measured markers required, alongside H&E, to accurately reconstruct large multiplex tissue imaging (MTI) panels while preserving biologically and clinically relevant information. miniMTI learns from paired same-section H&E–MTI data using a unified multimodal generative model that can condition on arbitrary combinations of measured marker channels, coupled with an iterative panel selection strategy to rank informative molecular anchors. Across colorectal and prostate cancer cohorts spanning two MTI platforms and over 40 million cells, miniMTI reduces a 40-marker MTI assay to H&E plus as few as three measured molecular markers, while accurately recovering withheld markers, preserving cellular phenotypes and spatial tissue architecture, and disease-associated molecular programs, including Gleason grade-linked signatures. By integrating histology context with sparse molecular grounding, miniMTI overcomes the limitations of morphology-only virtual staining and provides a scalable, cost-effective approach for expanding MTI-level biomarker coverage with retained biological interpretability and clinical relevance.

## Introduction

Multiplex tissue imaging (MTI) enables spatially resolved measurement of dozens of molecular markers within intact tissue sections, providing direct access to cellular states, tissue architecture, and microenvironmental organization. Rapid advances in MTI platforms such as CyCIF^1^, CODEX^2^, and mIHC^3^, together with their adoption in large-scale atlas efforts including the Human Biomolecular Atlas Program (HuBMAP)^4^ and the Human Tumor Atlas Network (HTAN)^4,5^, have established MTI as a cornerstone technology for spatial biology. However, high-plex antibody panels remain costly, time-intensive, and technically complex, limiting scalability and routine deployment across large cohorts and clinical workflows^6–9^.

A central challenge in MTI is balancing molecular coverage with practical constraints. Expanding panel size improves biological resolution but amplifies reagent consumption, staining cycles, imaging time, and cumulative tissue damage. Computational panel reduction and marker imputation methods partially address this trade-off by inferring unmeasured markers from a measured subset using models trained on fully multiplexed reference data^10–13^. While effective in reducing experimental burden, these methods still require non-trivial multiplex acquisition to provide sufficient molecular grounding that can capture the information content available, and they do not fundamentally alter the cost and complexity of high-plex spatial profiling.

In parallel, virtual staining approaches have sought to predict immunofluorescence (IF) or MTI marker expression directly from hematoxylin and eosin (H&E)–stained histology^9,14–21^ or immunohistochemistry (IHC) images^22^. Although attractive due to the low cost and ubiquity of H&E, morphology provides only indirect access to molecular state. As a result, H&E-only virtual staining exhibits strong performance for markers with clear morphological correlates but remains unreliable for many functional and signaling processes as well as target recognition in malignant cells, or immune or cancer associated fibroblast states at single-cell resolution. This limitation is not merely due to current model representation or learning capacity but reflects the inherent difficulty of modeling complex molecular states with tissue morphology as the sole input, rendering morphology-only inference unreliable for broad marker panels. This limits the ability to acquire biologically and clinically relevant information from H&E analysis.

Rather than serving as a substitute for molecular imaging, histology and MTI encode complementary biological information^23^. Effective integration of these modalities therefore requires models that can jointly leverage dense morphological context and direct molecular measurements. Crucially, spatial correspondence at the single-cell level is essential: approaches trained on paired, same-section H&E and MTI data consistently outperform those relying on adjacent sections, where cell-level misalignment and biological variability degrade molecular inference^24–26^. These observations suggest that accurate and scalable virtual multiplexing will require deliberate molecular anchoring rather than reliance on morphology alone.

Here, we introduce miniMTI, a unified framework for H&E-informed expansion of MTI using minimal molecular anchors. miniMTI is guided by a design principle: dense histologic context should be combined with a deliberately minimal set of experimentally measured markers to anchor inference of the remaining panel. H&E provides rich structural, morphological and contextual information, while sparse molecular measurements supply direct molecular specificity that resolves ambiguities inherent to morphology-only prediction. By learning from paired same-section H&E–MTI data, miniMTI unifies H&E-to-marker prediction and MTI-to-MTI imputation within a single generative model, and couples this with iterative panel selection to identify the most informative as well as cost effective and technically amenable anchors.

Using colorectal and prostate cancer cohorts profiled on distinct MTI platforms (Fig. 1a), we show that H&E combined with as few as three experimentally measured markers is sufficient to accurately reconstruct a 40-plex MTI panel while preserving biologically meaningful spatial organization with a high information content. Together these results establish miniMTI as a general, scalable strategy for expanding MTI-level biomarker coverage through histology-anchored diagonal integration of minimal molecular measurements.

**Figure 1.**
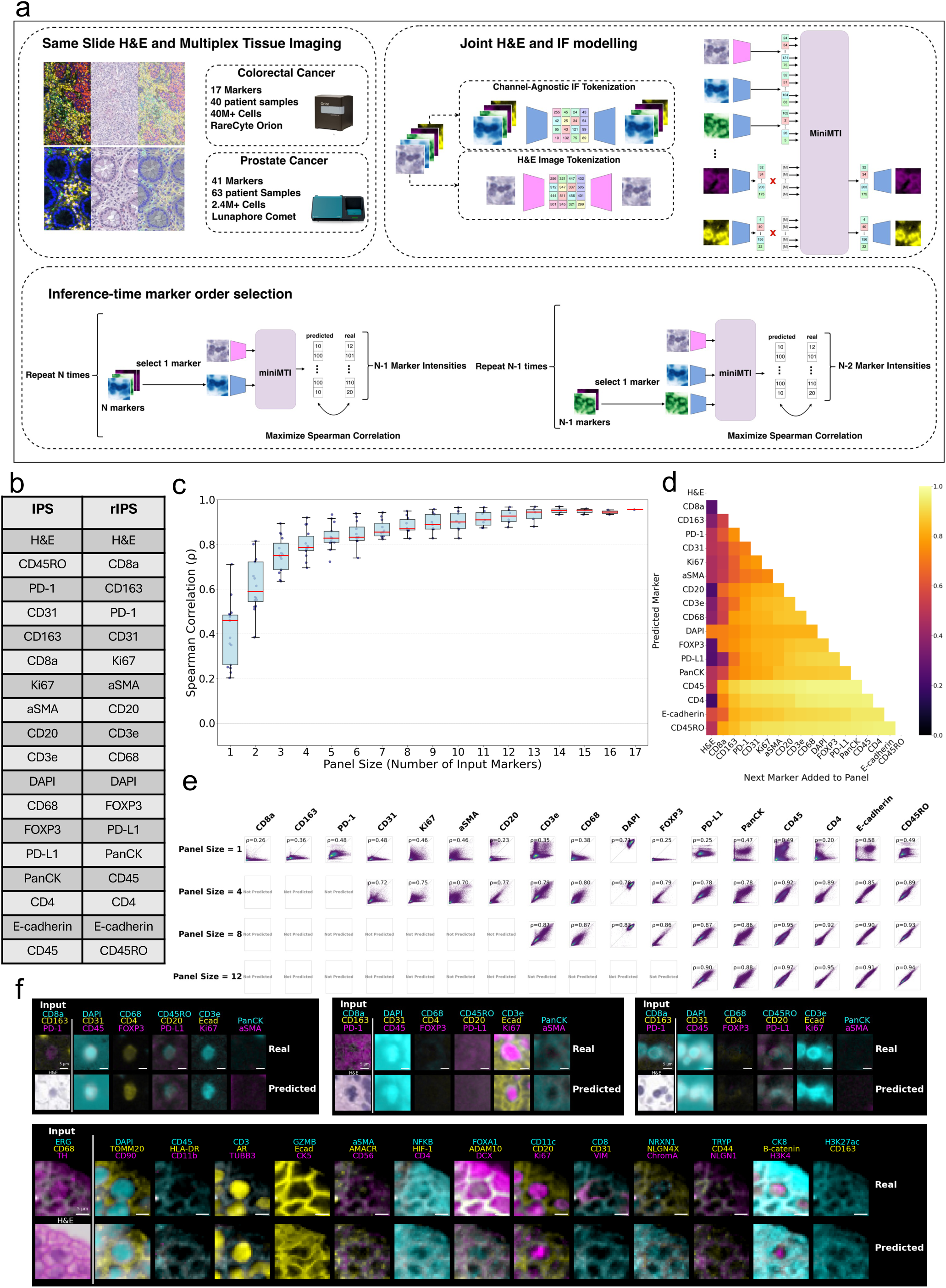
A generative masked language modelling approach for computational reduction of multiplex tissue imaging panels: a, Overview of dataset and model architecture. Paired H&E and multiplex IF images were collected on the same tissue sections for colorectal and prostate cancer cohorts using RareCyte Orion and Lunaphore Comet platforms, respectively. A generative masked language model was jointly trained on tokenized H&E and IF representations. After training, the model was probed to determine optimal subsets of markers that may be used to generate additional markers. b, Optimal marker orderings identified using IPS and rIPS algorithms for the 17-marker colorectal cancer panel. c, Boxplot showing improvement in prediction performance (Spearman correlation, ρ) for generated markers as a function of the number of input markers. d, Heatmap illustrating marker-wise impact of sequentially adding markers to the input panel. e, Scatter plots of real versus predicted marker intensities at the single-cell level. f, Representative examples of generated single-cell images using H&E and three IF markers as input. The top three examples are from the colorectal cancer cohort; the bottom example is from the prostate cancer cohort.

## Results

### miniMTI Overview

To enable flexible integration of histology with sparse molecular measurements, we designed miniMTI as a unified multimodal generative modelling framework^27–29^ that conditions on arbitrary combinations of H&E and IF marker channels while predicting the remaining withheld channels and retaining information content available from IF marker panels. Unlike prior virtual staining approaches that train marker-specific predictors or rely on H&E only, miniMTI is explicitly formulated to support flexible conditioning on variable marker subsets, allowing a single model to address both H&E-to-marker prediction and MTI-to-MTI imputation within the same architecture.

miniMTI represents both H&E and IF images as sequences of discrete visual tokens. Following prior work^29,30^, we employed vector-quantized generative adversarial networks (VQGAN)^31^ to learn compact, discrete codebooks for each modality, enabling high-fidelity compression of image patches into low-dimensional latent representations (Fig. 1a) (Methods). Separate tokenizers were trained for H&E and IF channels, with IF channels encoded independently using a shared IF tokenizer to allow marker-agnostic handling of variable panel compositions. The resulting token sequences were then used to train a bidirectional transformer with a masked language modeling objective^32,33^.

During training, entire image channels, either all tokens corresponding to an IF marker or the full RGB H&E images, were randomly masked and reconstructed. This channel-wise masking strategy enforces learning of cross-modal and cross-marker dependencies, enabling the model to infer missing molecular information from both histologic context and co-expression structure among measured markers. Importantly, masking was applied dynamically at every training iteration using newly sampled random subsets of channels, ensuring that the model learns to operate under arbitrary input configurations rather than overfitting to a fixed panel design.

At inference time, the model was provided with an H&E image together with a selected subset of experimentally measured IF channels. All remaining channels were masked and predicted by the model, yielding full-resolution synthetic marker images. This formulation naturally supports virtual staining (H&E→IF), marker imputation (IF→IF), and hybrid conditioning (H&E+IF→IF) within a single framework, and enables a direct interrogation of how prediction fidelity varies as a function of input panel composition.

After training a single unified miniMTI model, we applied an iterative panel selection (IPS) strategy^34^ that ranks IF markers based on their marginal contribution to reconstructing the remaining panel. Starting from H&E, IF markers were added sequentially according to the improvement in overall prediction accuracy, yielding an ordered list in which early IF markers define an optimized subset of size *k* (Methods). We additionally evaluated a complementary heuristic, reverse IPS (rIPS)^29^ heuristic that constructs the ordering in reverse by prioritizing IF markers that were easiest to predict from the remaining set. In our experiments, rIPS provided slightly improved performance and is used for subsequent analyses.

Figure 1b shows the IPS and rIPS ordering for a 17-marker colorectal cancer (CRC) panel^23^ acquired on the RareCyte Orion imaging platform, together with the progressive improvement in prediction performance as additional anchors are incorporated (Figs. 1c-e). We quantified performance using single-cell mean intensities computed within the segmented cell boundary of the central cell in each crop and evaluated agreement with ground truth provided by the complete IF set using Spearman correlation across cells. While our primary analyses focused on cell-level quantitative features to align with standard MTI workflows^35,36^, there is growing interest in complementing or replacing classical single-cell features with deep learning-derived image representations that better capture morphological context and reduce sensitivity to segmentation error^37,38^. Consistent with this trend, the generative nature of miniMTI enables synthesis of spatially coherent marker images (Fig. 1f, Supp. Fig. 1), preserving fine-grained tissue structure beyond summary intensity metrics.

### miniMTI outperforms late-fusion integration of single-modality foundation models

We next evaluated whether the advantages of miniMTI arise from its unified multimodal formulation or could instead be matched by late-fusion integration of state-of-the-art single-modality foundation models for histology and multiplex imaging. As a representative late-fusion baseline, we combined UNI^39^, a general-purpose histopathology foundation model trained on large-scale H&E data, with KRONOS^40^, a spatial proteomics foundation model pre-trained on MTI. This design reflects a common strategy in multimodal learning in which independently trained encoders are fused at the feature level to perform downstream prediction.

For this benchmark, we formulated a regression task in which the input consists of H&E images together with the top three rIPS-selected markers (CD8a, CD163, and PD-1), and outputs were the mean expression values of the remaining fourteen markers (Methods). For miniMTI, we generated full-resolution predictions for the withheld channels and computed mean pixel intensities within the segmented cell boundaries. For the UNI+KRONOS late-fusion baseline, we extracted UNI and KRONOS feature embeddings from the same inputs, concatenated features, and trained a multilayer perceptron to directly predict the fourteen mean intensities.

Across markers, miniMTI consistently outperformed the UNI+KRONOS late-fusion model (Fig. 2a), with the performance gains reproducible across nine held-out patient samples (Fig. 2b) Supplementary Fig. 2 reports Pearson correlation coefficients). We further compared UNI-only and KRONOS-only baselines, as well as miniMTI variants with and without H&E input (Methods). In all settings, early multimodal fusion in miniMTI provided a systematic advantage over single-modality models and late-fusion integration.

**Figure 2.**
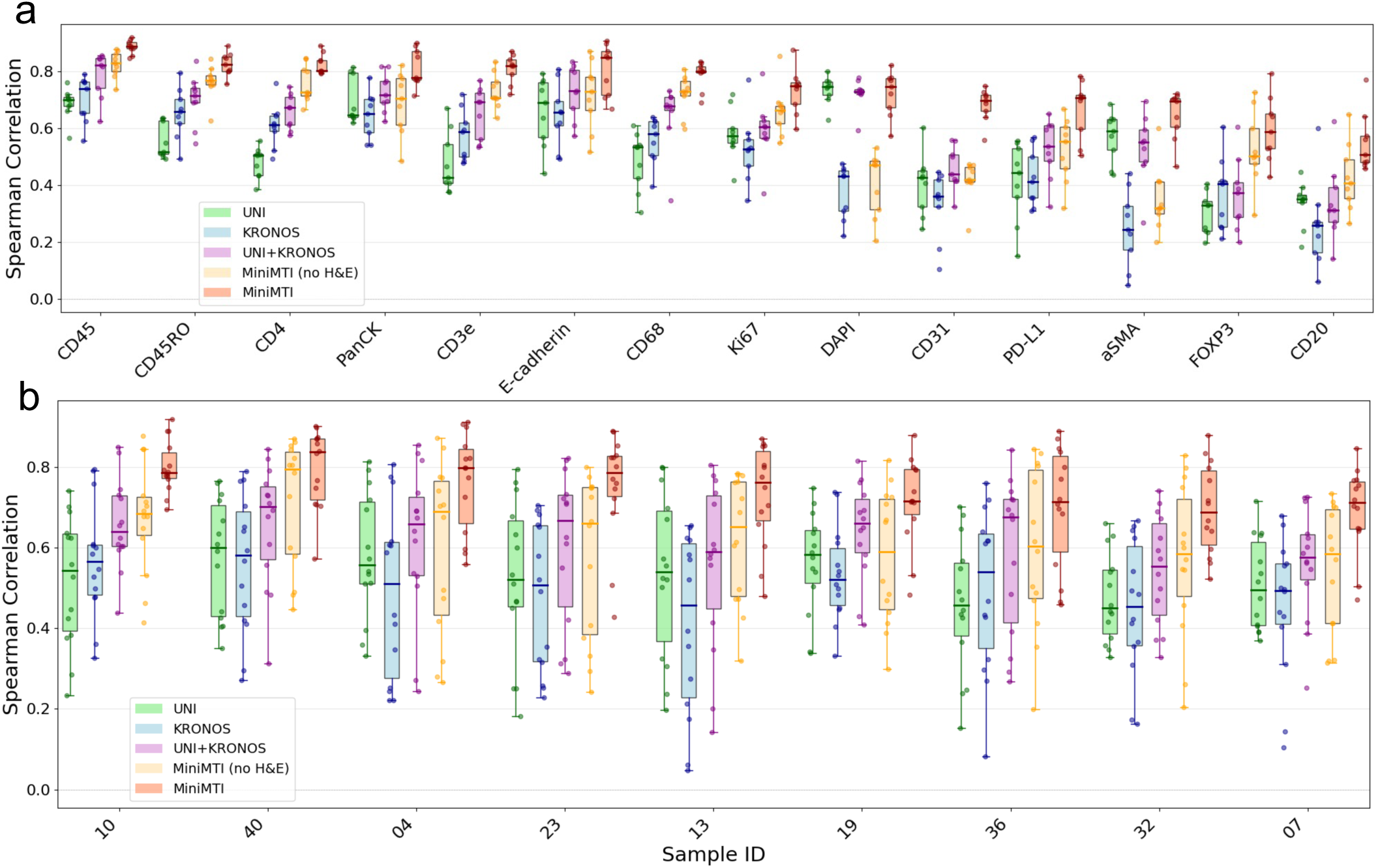
miniMTI outperforms state-of-the-art foundation models for protein expression prediction. **a**, Marker-wise comparison of prediction performance (Spearman correlation, ρ) across models, including foundational model ensembles (UNI, KRONOS, and UNI+KRONOS) and miniMTI configurations using IF-only inputs, and IF plus H&E inputs. **b**, Sample-wise comparison of the same models across nine held-out colorectal cancer patient samples. miniMTI consistently achieves higher correlation across markers and samples, with additional gains when incorporating H&E.

These results highlight a key methodological distinction: in late-fusion approaches, cross-modal relationships are learned only in a shallow, task-specific predictor layered on top of fixed unimodal representations. In contrast, miniMTI learns joint representations of histology and molecular imaging through channel-wise masked modeling, enabling cross-modal dependencies to be captured throughout the depth of the network. This early fusion allows miniMTI to exploit both co-expression structure among markers and morphological context simultaneously, rather than combining modalities only after independent encoding. Together, these findings demonstrate that sparse molecular anchoring within a unified multimodal generative framework is fundamentally more effective than post hoc fusion of single-modality foundation models for virtual marker prediction.

### Leveraging H&E to reduce high-plex biomarker panels

To disentangle the respective contributions of histology and sparse molecular input to marker reconstruction, we evaluated miniMTI under three input conditions: H&E alone, three-marker input identified by rIPS (CD8a, CD163, and PD-1) alone, and the combined H&E plus three-marker condition. This experimental design enables direct quantification of incremental predictive value contributed by each modality and isolates the extent to which sparse molecular anchoring mitigates the intrinsic limitation of morphology-only inference.

Across nine CRC test samples, combining H&E with three measured markers consistently improved prediction accuracy for all withheld channels relative to either modality alone (Fig. 3a). The combined condition outperformed H&E-only prediction for all markers and exceeded the three-marker-only condition, demonstrating that histologic context and molecular anchors provide complementary and non-redundant information.

**Figure 3.**
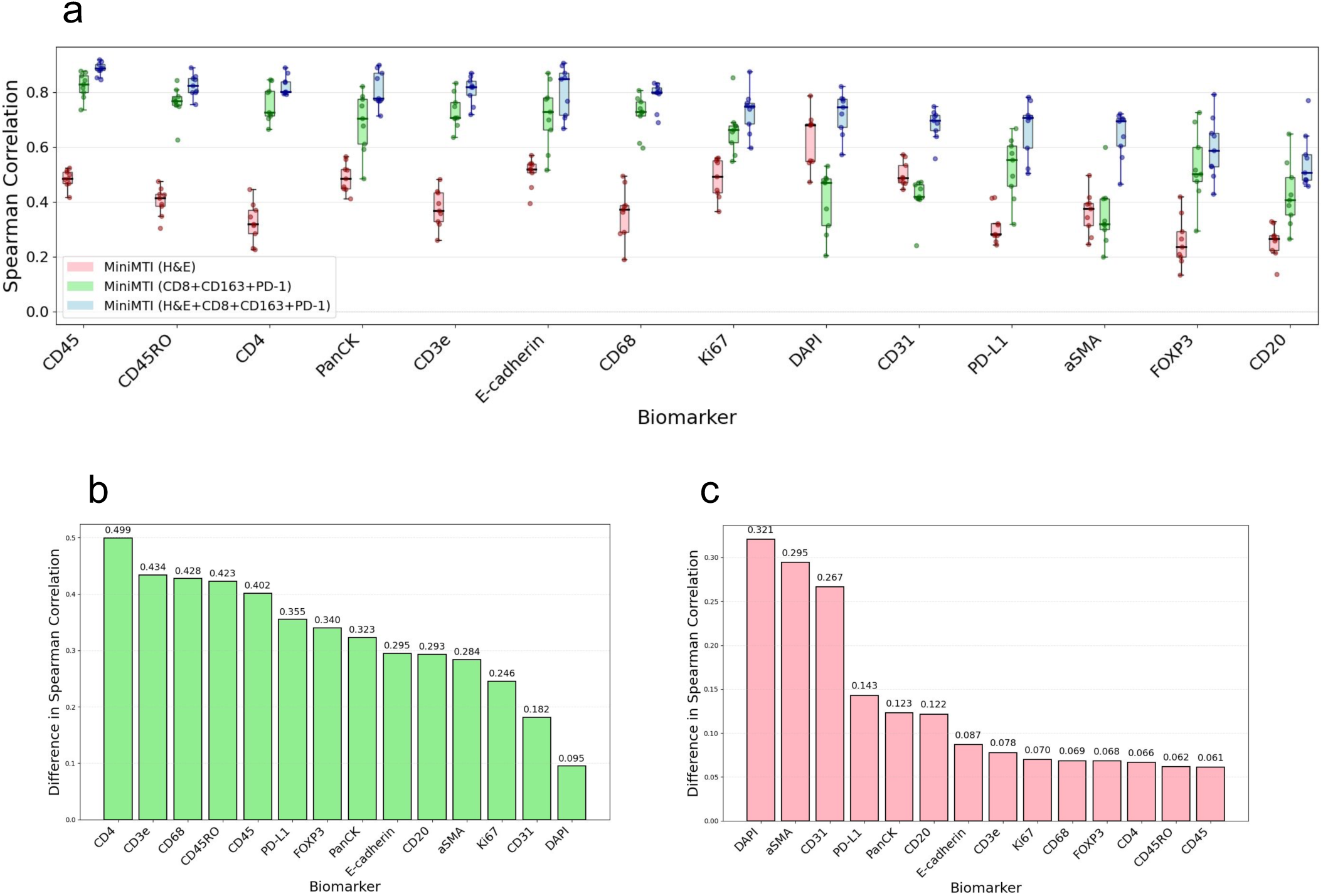
Quantifying the complementary contribution of H&E and MTI to predictive performance. a, Marker-wise comparison of prediction performance (Spearman correlation, ρ) across miniMTI input configurations: H&E only, IF only (CD8+CD163+PD-1), and combined IF plus H&E. b, Per-marker gain in prediction performance when three IF markers are added to H&E input. c, Per-marker gain in prediction performance when H&E is added to three IF markers as input.

To quantify the benefit of sparse molecular anchoring beyond morphology, we computed the per-marker change in Spearman correlation between H&E+3 marker and H&E-only conditions (Fig. 3b). Broad improvements were observed across the panel, with the largest gains observed for immune markers, consistent with anchor-driven learning of co-expression and mutual exclusion relationships. For example, prediction accuracy for CD3e and CD4 increased substantially when conditioning on CD8a, reflecting recovery of biologically expected T cell subtype structure that is weakly encoded in morphology alone.

Conversely, to quantify the contribution of histological context beyond molecular input, we computed the difference between H&E+three marker and three-marker-only conditions (Fig. 3c). Incorporating H&E produced strong gains for markers with clear morphologic correlates, including DAPI (nuclear content), αSMA (stromal structures), and CD31 (vasculature), and improved performance across the full panel. These results indicate that histology provides valuable spatial and structural context that cannot be recovered from sparse molecular measurement alone.

Together, these analyses demonstrate the predictive gains achieved by miniMTI are not attributable to either modality in isolation. Instead, accurate reconstruction of high-plex panels emerges from the deliberate integration of dense histologic context with sparse molecular grounding. These findings are consistent with our benchmarking results in Fig. 2a, where the combined UNI+KRONOS late-fusion model outperforms the KRONOS-only model alone across all markers. Together, these analyses demonstrate that histological information contributes positively to marker prediction performance and that its value is maximized when integrated with sparse molecular measurements in a multimodal framework. This complementary contribution explains why miniMTI consistently outperforms H&E-only virtual staining and IF-only imputation and supports the design principle that minimal molecular anchoring is required to overcome the marker-dependent performance ceiling of morphology-only inference.

### miniMTI recapitulates spatial patterns from ground truth data

A key advantage of MTI is access to spatially resolved marker expression at single-cell resolution. We therefore evaluated whether miniMTI preserves higher-order tissue organization beyond marker-wise accuracy by assessing the concordance of spatial pattern between ground-truth and imputed data. To this end, we performed CellCharter^41^ spatial clustering on both experimentally measured and miniMTI-imputed marker profiles across nine CRC samples (Methods).

CellCharter consistently identified reproducible spatial niches corresponding to major tissue compartments, including tumor epithelium, stromal regions, immune infiltrates, and vascular structures (Fig. 4). Spatial clustering assignments derived from imputed data show substantial concordance with ground truth clusters derived from the experimentally measured MTI marker intensities, with adjusted Rand index (ARI) values ranging from 0.3 to 0.41 across samples (Fig. 4a). Visual comparison confirmed that miniMTI recovered the spatial distribution of major tissue compartments (Fig. 4b). For instance, in sample CRC04, both ground-truth and imputed data identified a prominent αSMA-enriched stromal compartment and a distinct immune-infiltrated region characterized by high CD45, CD3e, and CD8 expression. Detailed results for individual tissue compartments across all samples are provided in Supp. Fig. 3.

**Figure 4.**
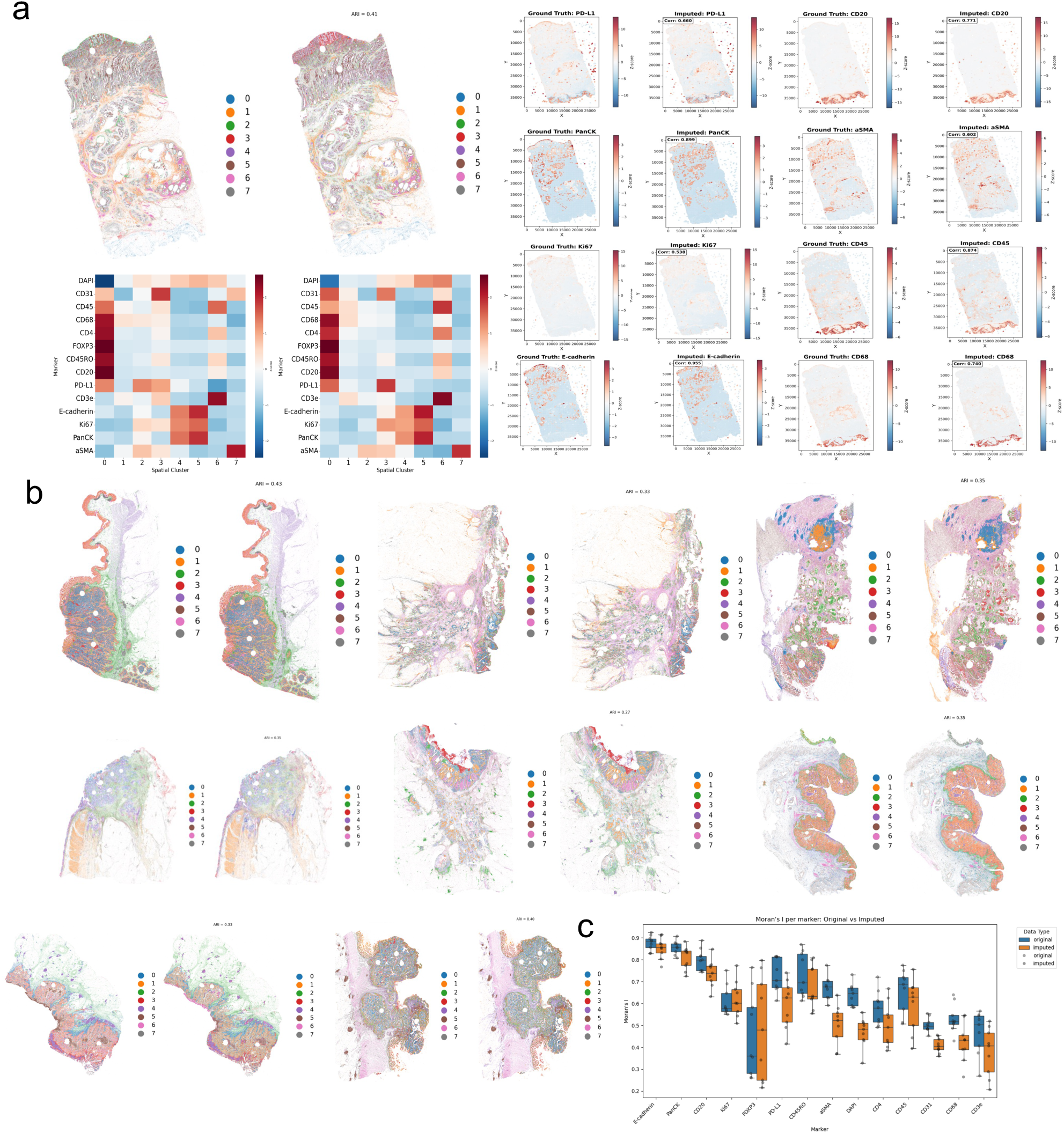
Spatial reconstruction of MTI using H&E and minimal anchor markers. **a**, Spatial clustering maps derived from ground truth MTI (left) and miniMTI-predicted MTI using H&E plus 3 markers (right). Each color denotes a distinct spatial state or niche defined by CellCharter, showing close agreement in tissue architecture and regional organization between measured and imputed data (right panel). **b**, Spatial clustering maps for different tissue samples, comparing ground truth MTI (left) and imputed MTI (Right). Consistent spatial patterns and niche organization were preserved across tissues, demonstrating robust generalization of the model. **c**, Comparison of spatial autocorrelation (Moran’s I) for each marker between ground truth (blue) and imputed (orange) data. Box plots summarize distributions across samples, indicating that the model preserves marker-specific spatial patterning.

To assess molecular consistency within spatial niches, we compared cluster-level marker expression profiles between ground-truth and imputed data. Cluster-level marker expression heatmaps demonstrated strong concordance between matched spatial niches, with Pearson correlation coefficients for cluster centroids ranging from r = 0.67 to 0.91 (P < 0.001).

Marker-specific spatial expression maps further confirmed preservation of fine-grained spatial patterns for both immune and structural markers (Fig. 4c). Immune markers, including CD3e, CD8, and PD-L1, exhibited high spatial correspondence between ground-truth and imputed data (r > 0.91), recapitulating immune enrichment at the tumor-stroma interfaces and exclusion from tumor epithelial regions. Structural markers such as E-cadherin and PanCK similarly retained their characteristic epithelial localization patterns. Across spatial niches, stromal and epithelial compartments showed the highest concordance between ground-truth and imputed data, while spatially heterogeneous immune subpopulations exhibited slightly lower but still significant agreement.

We next quantified preservation of global spatial organization using Moran’s I for each marker. Across CRC samples, original MTI measurements showed slightly higher global spatial autocorrelation than imputed values. On average, Moran’s I was approximately 0.07 higher in the original data. Although statistically significant after accounting for sample-level variability, this difference is modest, indicating that miniMTI introduces a mild spatial smoothing while largely preserving the global spatial organization of biomarker expression.

Together, these analyses demonstrate that miniMTI preserves biologically meaningful tissue architecture and higher-order spatial organization, not merely marginal marker distributions. The ability to recover spatial niches and coordinated expression patterns from minimal molecular input highlights the importance of joint multimodal modeling for faithful reconstruction of tissue microenvironments.

### Extended validation in prostate cancer

To assess the generalizability of miniMTI beyond colorectal cancer and across distinct MTI technologies, we applied the framework to an independent in-house cohort of 63 prostate cancer core needle biopsies profiled with 41 IF markers using the Lunaphore COMET platform, together with same-section H&E. This cohort differs substantially from the CRC dataset in tissue architecture, disease biology, and staining chemistry, providing a stringent test of model robustness.

As in the CRC analysis, marker-wise prediction accuracy increased systematically with panel size. Notably, this trend was preserved in the prostate cohort despite the substantially larger 41-marker panel, demonstrating that miniMTI scales effectively to higher-plex MTI. Across the full marker set, H&E combined with 3 selected markers (CD68, TH, ERG) outperforms H&E-only prediction across most markers, with additional gains observed for H&E+6 (CD68, TH, ERG, NLGN1, CK8, GZMB) and H&E+9 (CD68, TH, ERG, NLGN1, CK8, DAPI, GZMB, CD4, CK5) configurations (Fig. 5a). These improvements are consistent across Gleason grades, indicating stable performance across disease states, varying levels of histopathological heterogeneity and clinical relevance (Fig. 5b).

**Figure 5.**
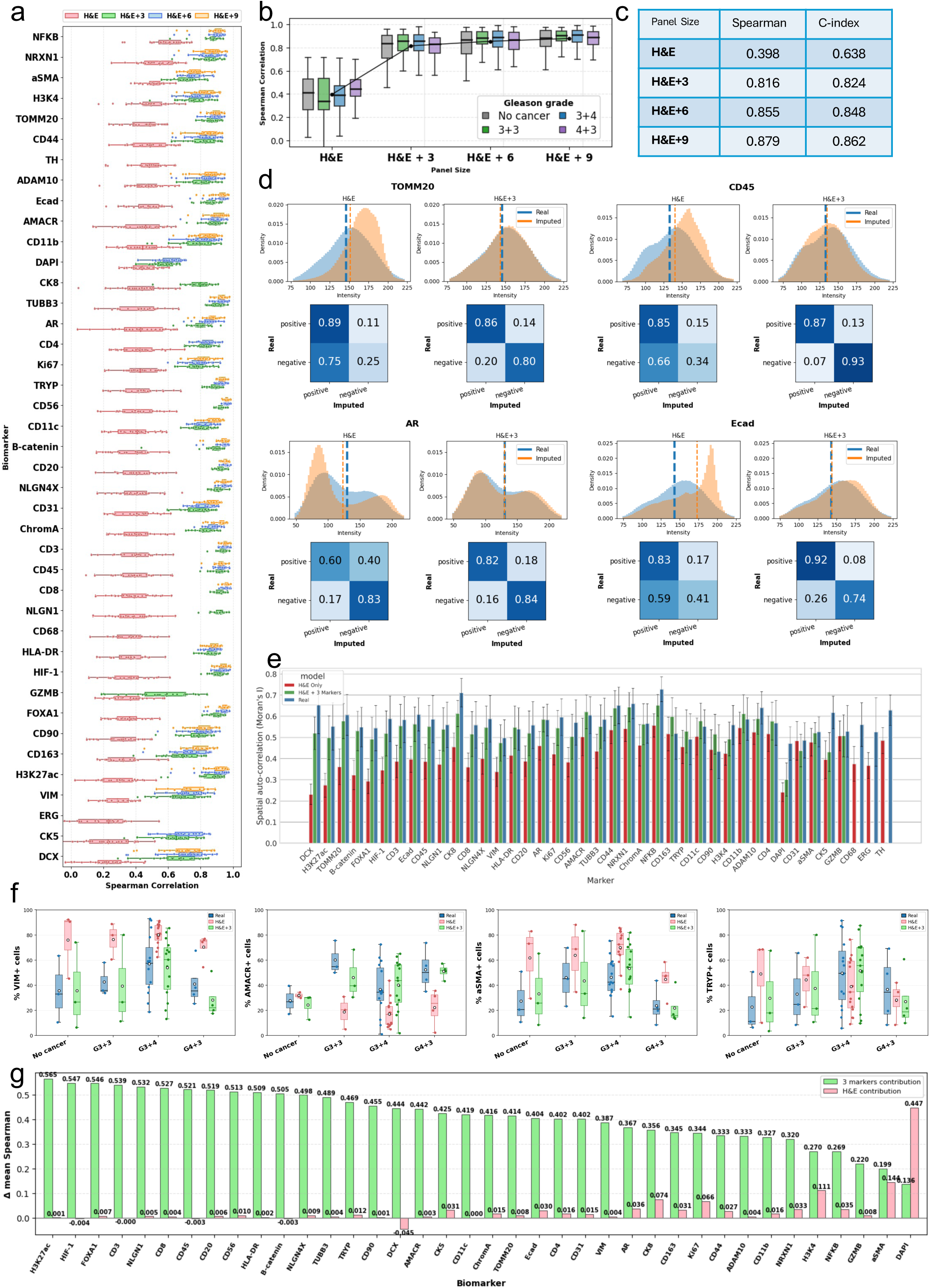
Extension to a 40-marker prostate cancer dataset demonstrates scalability and biological fidelity. a, Per-marker prediction accuracy (Spearman correlation between real and imputed single-cell intensities) for four panel sizes: H&E-only, H&E+3, H&E+6, and H&E+9 anchor markers. b, Distribution of Spearman correlation stratified by Gleason grade (no cancer, 3+3, 3+4, 4+3) for each panel size. c, Summary table reporting aggregate Spearman correlation and C-index for each panel size. d, Representative markers (TOMM20, CD45, AR, and E-cadherin) showing recovery of expression distributions. Overlaid histograms show real (blue) and imputed (orange) intensities for H&E-only and H&E+3. Dashed lines indicate two-component GMM decision boundaries derived from real data (bold blue) and imputed data (orange dashed line). For the gating analysis, positivity thresholds were defined using the real-derived gate only and then applied to imputed data. Confusion matrices (rows = real, columns = imputed) summarize agreement of positive/negative labels, highlighting higher concordance and reduced false positives with H&E+3. e, Marker-wise recovery of spatial organization quantified by spatial autocorrelation (Moran’s I), comparing real measurements to imputed values (H&E-only versus H&E+3 markers). f, Preservation of disease-associated trends. Fraction of marker-positive cells across Gleason groups for progression-associated markers (VIM, AMACR, αSMA, Tryptase), comparing real with imputed predictions from H&E-only and H&E+3, using the same GMM-based gating as in subpanel d. g, Marker-wise decomposition of performance gains, showing the relative contributions from H&E morphology versus the three anchor markers to the mean Spearman improvement.

Aggregate performance metrics, including both Spearman correlation and concordance index (C-index), show monotonic improvement as additional anchors were added (Fig. 5c). Notably, miniMTI achieved a strong baseline performance even in the H&E+3 setting, with Spearman increasing from 0.398 (H&E) to 0.816 (H&E+3). In contrast to prior H&E-only virtual staining approaches that often rely on rank-based metrics while exhibiting compressed dynamic range^14,15^, miniMTI demonstrates substantial recovery of absolute intensity structure with minimal molecular input.

To assess whether sparse molecular anchoring preserves the full dynamic range and population structure of marker expression, we compared real and imputed intensity distributions for representative functionally relevant markers, including TOMM20, CD45, AR, and E-cadherin (Fig. 5d) (full markers in Supp. Fig. 4). Under H&E-only input, imputed intensities exhibited reduced dynamic range or distorted distribution, and poorer separation of positive and negative populations. In contrast, miniMTI with H&E+3 recapitulated the shape of the ground-truth distributions of the experimentally measured marker intensities, and restored clear population structure. Using a positivity threshold derived from the real staining data via two-component Gaussian mixture modeling (GMM) (Methods), miniMTI with H&E+3 accurately reproduced positive and negative population structure, with high agreement between real and imputed calls. For instance, for CD45, negative-population agreement increased from 0.34 (H&E) to 0.93 (H&E+3), reflecting a marked reduction in false positives.

We next evaluated preservation of global spatial organization using marker-wise Moran’s I (Fig. 5e). With H&E-only input, imputed marker values exhibited substantially weaker and more variable spatial autocorrelation compared to experimentally measured MTI marker intensities (ground truth), indicating inconsistent preservation of spatial structure. Incorporation of three experimentally measured markers alongside H&E markedly mitigated spatial signal loss, with the mean difference in Moran’s I between original and imputed values reduced to 0.033 (p<0.001). Compared to the H&E-only setting, the reduction in Moran’s I was markedly attenuated, demonstrating improved recovery of spatial patterns with minimal molecular input. Although statistically significant, this difference was small, indicating that most of the global spatial organization present in the original MTI data was preserved under minimal molecular input.

Finally, to assess preservation of clinically relevant trends, we investigated epithelial and stromal markers associated with disease progression (AMACR, αSMA, VIM, and Tryptase) across Gleason grade groups^42^. The imputed data recapitulated the same grade-dependent shifts in marker positivity observed in the ground-truth measurements (Fig. 5f), indicating miniMTI preserves clinically meaningful marker-disease relationships.

Quantitatively, per-sample positivity fractions show consistently stronger concordance between real and imputed data under H&E+3 condition compared to H&E-only across all four progression-associated markers. Specifically, correlations improve for VIM (r=0.536 vs. r=0.853), AMACR (r=−0.011 vs. r=0.625), αSMA (r=0.842 vs. r=0.933), and Tryptase (r=0.331 vs. r=0.871), indicating substantial gains in sample-level agreement with the addition of sparse molecular anchors that could influence both information context and clinical impact.

Consistent with these trends, β-distribution fitting (Methods) demonstrated that the H&E+3 condition yielded parameter estimates markedly closer to those derived from experimentally measured MTI staining (ground truth) than H&E-only (e.g., VIM: Real (17.27, 21.92) vs. H&E (118.55, 38.05) vs. H&E+3 (10.67, 16.48); Tryptase: Real (8.60, 15.67) vs. H&E (15.39, 23.07) vs. H&E+3 (9.56, 16.69); Supp. Fig. 5). These results indicate improved recovery of clinically meaningful, high information content, progression-linked patterns rather than distortion driven by H&E-only inference.

Consistent with our colorectal cancer analysis (Fig. 3), Figure 5g shows the same complementary and non-redundant contributions of histology and sparse molecular anchoring across biomarkers. For example, DAPI is largely morphology-driven (H&E contribution 0.447 versus 3-markers contribution 0.136), whereas αSMA benefits from both H&E and space molecular anchoring.

Together, these results demonstrate that the design principles underlying miniMTI generalize across cancer types, tissue architectures, and imaging platforms. This cross-context robustness supports miniMTI as a general framework for histology-anchored expansion of multiplex tissue imaging, rather than a dataset- or platform-specific solution.

### Confidence estimation using miniMTI-derived likelihood scores

A practical requirement for deploying virtual staining methods is the ability to assess prediction reliability. A key advantage of miniMTI’s language-modelling inspired framework is the ability to estimate prediction confidence directly from sequence likelihood. Specifically, each image channel is tokenized into a sequence of discrete visual tokens, and the model is trained to predict a probability distribution over the token vocabulary at each masked position. Following prior work on masked language model scoring^43^, we compute a pseudo-log likelihood (PLL) score for each tokenized image sequence by aggregating the log-probabilities assigned to the observed tokens. Higher PLL values indicate that the model assigns a higher likelihood to the input image under the learned data distribution, providing a quantitative, model-intrinsic measure of prediction confidence.

We first evaluated how confidence changes with additional molecular input by computing ground-truth PLL distributions of experimentally measured MTI marker images to predicted distributions generated from H&E-only and H&E+three-marker input panel (Fig. 6a).

**Figure 6.**
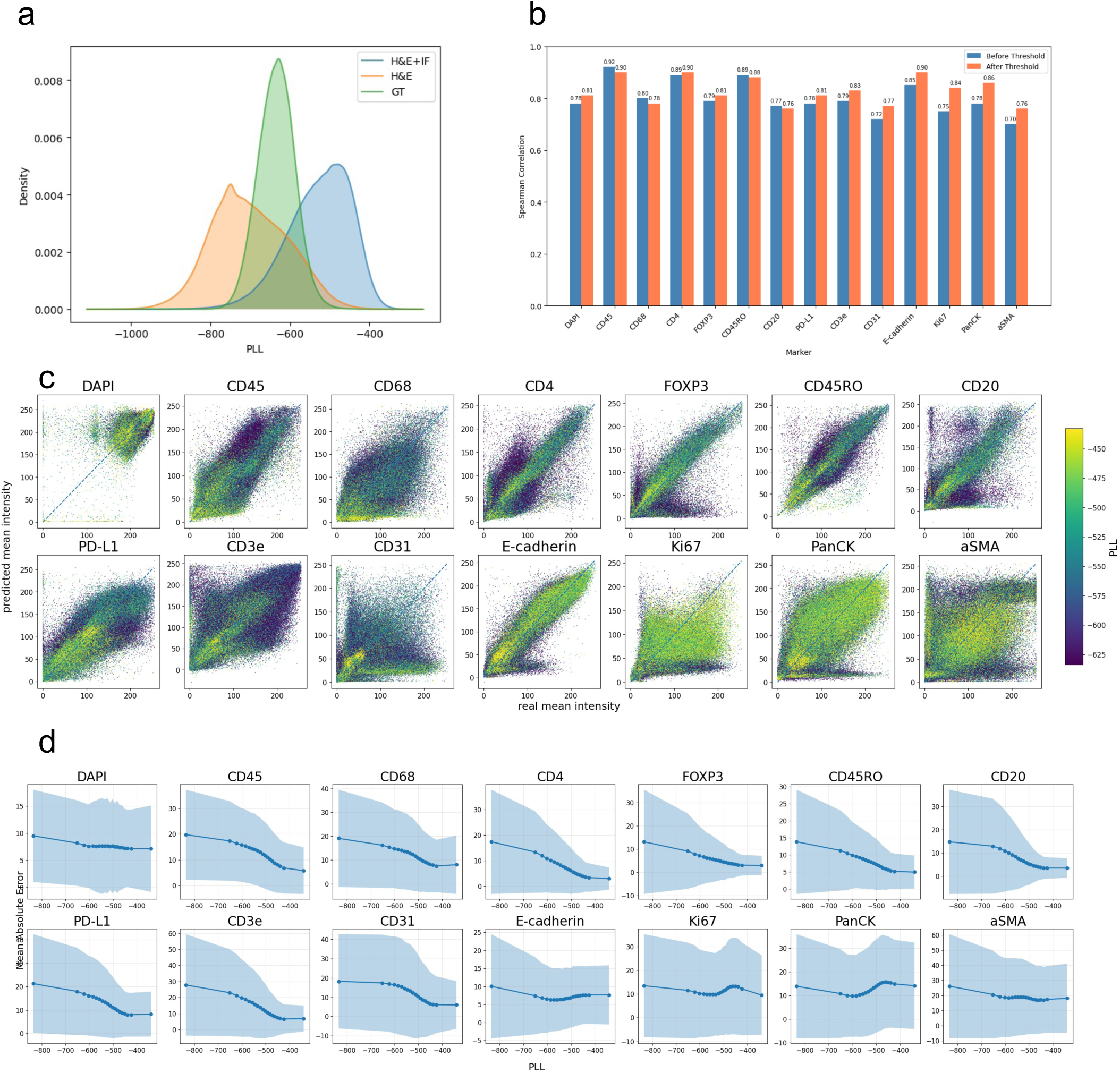
miniMTI provides model intrinsic confidence estimation. **a**, Density plot for pseudo log-likelihood (PLL) scores produced using the CRC test set with three different input conditions, ground truth images, images generated using H&E only as input, and images generated using H&E plus three additional IF markers (CD8+CD163+PD-1). b, Barplots depict prediction performance for 14 markers from the CRC test set generated using H&E plus three additional IF markers as input for the whole test set, and for the top 50% of cells with the highest PLL scores. **c**, Scatterplots show real versus predicted mean intensities for the 14 predicted CRC markers colored by PLL scores. **d**, Plot depicts mean absolute error for the 14 predicted mean intensities binned by PLL score.

Incorporation of the three experimentally measured markers shifts the PLL distribution upward, from the lower range of the ground-truth distribution toward its upper range, indicating increased model confidence when sparse molecular anchors were provided alongside morphology. In contrast, H&E-only predictions were enriched for low-PLL values, consistent with the intrinsic uncertainty of morphology-only inference.

We next evaluated whether PLL can be used to identify unreliable predictions. Retaining only the top 50% of predictions ranked by PLL increases per-marker Spearman correlations for 10 of 14 markers (Fig. 6b). Consistent with this, overlaying PLL scores on scatter plots of ground-truth versus predicted mean intensities revealed that low-PLL predictions were enriched among outliers (Fig. 6c).

Finally, we examined the relationship between confidence and accuracy by plotting mean absolute error (MAE) as a function of PLL. For most markers, higher PLL was associated with progressively lower absolute error between predicted and ground-truth intensities (Fig. 6d), demonstrating that miniMTI-derived likelihood scores provide a meaningful and practical estimate of prediction reliability. Together, these results demonstrate that miniMTI provides a model-intrinsic, calibration-free measure of prediction confidence that can be used to flag uncertain virtual stains and guide downstream analysis by allowing focus on high quality data..

## Discussion

In this study, we present miniMTI, a general framework for histology-anchored expansion of multiplex tissue imaging using minimal molecular anchors. By integrating routine H&E with minimal multiplex immunofluorescence imaging, miniMTI addresses a fundamental limitation of morphology-only virtual staining: the intrinsic information bottleneck that constrains reliable inference of many functional and immune biomarkers from histology alone. Rather than treating histology as a surrogate for molecular imaging, miniMTI explicitly leverages the complementary information encoded in tissue morphology and sparse molecular measurements to enable accurate reconstruction of high-plex marker panels.

Our results demonstrate that modern generative modeling approaches can be effectively applied to multiplex tissue imaging to expand molecular coverage while reducing experimental burden. Masked language models have shown strong performance across domains, including natural language^44^, computer vision^33^, and protein modeling^45^, and have increasingly been adopted for unified multimodal representation learning^46,47^. Building on these advances, miniMTI leverages a natively multimodal architecture to jointly model histology and molecular imaging data, enabling the generation of high-fidelity synthetic marker channels from limited experimental input.

A central insight from our work is that sparse molecular anchoring is not merely additive but structurally transformative. Even a small number of experimentally measured markers fundamentally alters the identifiability of many biomarkers that are weakly or ambiguously encoded in morphology, recovering both marker-specific expression patterns and higher-order spatial organization. This design principle explains why miniMTI consistently outperforms H&E-only virtual staining, IF-only imputation, and late-fusion integration of single-modality foundation models, and why its performance scales monotonically with increasing panel size across cancer types and imaging platforms.

Importantly, miniMTI is designed for practical utility, not only improved prediction accuracy. By reducing the number of experimentally required antibodies to a minimal anchor set, miniMTI can substantially lower reagent cost, imaging time, and assay complexity while retaining high information content through recovery of the remaining panel. This shift expands the range of feasible acquisition platforms: instead of relying on highly multiplexed workflows (e.g., CyCIF/CODEX), anchor acquisition can be performed using standard low-plex immunofluorescence available in most laboratories. In particular, an anchor set of three IF markers is compatible with routine 4-color IF workflows (with DAPI for nuclear segmentation) and with emerging 5-color clinical-grade platforms. This makes histology-anchored virtual multiplexing accessible on systems that measure a limited number of analytes per section, such as Orion^23^ and Leica-based^48^ workflows, while still enabling downstream analyses that typically require higher-plex MTI (e.g., cell phenotypes, spatial neighborhoods, and clinically relevant tissue states).

While our study focuses on protein-based multiplex imaging, the general framework is conceptually extensible to other high-dimensional, cost-prohibitive spatial assays such as spatial transcriptomics. However, important challenges arise when scaling from panels of tens of protein markers to hundreds or thousands of gene targets. In particular, our iterative panel selection strategy becomes computationally intractable as the search space grows combinatorially with panel size. This limitation highlights the need for more scalable optimization strategies. One promising direction is the use of reinforcement learning-based approaches, which showed improved efficiency and solution quality over exhaustive or greedy search in related high-dimensional selection problems^49^. Incorporating such strategies could enable the practical extension of miniMTI to substantially larger molecular panels.

More broadly, miniMTI illustrates a path toward reducing the experimental and logistical barriers associated with high-plex spatial profiling through histology-anchored diagonal integration of minimal molecular measurements^50^. By combining minimal molecular measurements with histology-driven inference, this approach reduces experimental cost and technical complexity while preserving, and in many cases expanding, the effective information content available from each tissue section, thereby increasing the practical utility of spatial biomarker analysis in both research and clinical environments. Future work will focus on extending miniMTI to additional tissues, disease contexts, and imaging platforms, and refining model efficiency and robustness to support broader deployment in translational research settings.

## Acknowledgements

This work was carried out with major support from the National Cancer Institute (NCI) Human Tumor Atlas Network (HTAN) Research Centers at OHSU (U2CCA233280), and funding from the Cancer Early Detection Advanced Research Center at Oregon Health & Science University, Knight Cancer Institute (CEDAR 2023-1796). Y.H.C. is supported by R01 CA253860, R01 CA276224, U01 294548, and the Kuni Foundation Imagination Grants. S.E.E. is supported by Prostate Cancer Foundation (GCNCR1865A) and International Alliance for Cancer Early Detection (ACED 2022-1508). The research reported in this publication used computational infrastructure supported by the Office of Research Infrastructure Programs, Office of the Director, National Institutes of Health, under Award Number S10OD034224. The content is solely the responsibility of the authors and does not necessarily represent the official views of the National Institutes of Health.

## Methods

### Multiplex Tissue Imaging of Colorectal Cancer Samples

Multiplex tissue imaging data for colorectal cancer were obtained from the publicly available CRC-ORION cohort generated by Lin *et al.* (Nature Cancer, 2023)^23^. This dataset comprises same-section paired H&E and high-plex immunofluorescence (IF) images acquired using the Orion platform, enabling precise cell-level correspondence between histology and molecular markers. The panel includes 17 protein markers spanning epithelial, immune, stromal, and functional markers, including Hoechst, CD31, CD45, CD68, CD4, FOXP3, CD45RO, CD20, PD-L1, CD3e, E-cadherin, Ki67, PanCK, and αSMA. Images were downloaded via the public Human Tumor Atlas Network data portal. No additional staining or imaging was performed. This dataset was used for all colorectal cancer analyses in this study.

### Human prostate cancer samples

Prostate tumor specimens analyzed in this study were obtained from treatment-naïve patients undergoing diagnostic needle biopsy at Oregon Health & Science University (OHSU) and University College London (UCL). Tissue collection was performed following written informed consent and in accordance with institutional review board–approved protocols at both sites. Use of human specimens was approved by the OHSU IRB (IRB#4918) and the UCL Biobank (Project NC27.22). All associated clinical and pathological data were de-identified prior to downstream analyses.

### Multiplex Tissue Imaging of Prostate Cancer Needle Biopsy Samples

Slides were deparaffinized and antigen retrieved with PT Module (Epredia Lab Vision^TM^) using pH 9 Dewax and HIER buffer H (TA999-DHBH, Epredia) at 102°C for 60 min. Then, slides were rinsed with 1X PBS and loaded into the COMET MK03 Imaging Chip. Imaging reagents were loaded into Lunaphore COMET (Lunaphore Technologies) to perform automated multiplex tissue immunofluorescence imaging (MTI). Following primary mouse and rabbit antibodies were used (two per cycle) along with DAPI (Thermo Scientific, cat no: 62248, 1/1000 dilution): AR (D6F11,1:100), ADAM10 (EPR5622, 1:100), AMACR (bs-0840R), aSMA (1A4, 1:3000), β-catenin (14RUO, 1:100), CD56 (VIN-IS-53, 1:100), CD68 (KP1, 1:50), CHGA (EP1030Y, 1:100), CK5 (EP1601Y, 1:200), CK8 (EP1628Y, 1:400), CD90 (EPR3133, 1:500), CD45 (2B11+PDy/26, 1:100), CD11b (EPR1344, 1:200), CD3 (UCHT1, 1:100), CD4 (EPR6855, 1:200), CD11c (EP1347Y, 1:500), CD20 (L26, 1:100), CD31 (EP3095, 1:400), CD8 (4B11, 1:50), CD44 (EPR1013Y, 1:500), CD163 (6E7A9, 1:100), DCX (2G5, 1:50), ECAD (EPY700Y, 1:800), ERG (EPR3864, 1:100), FOXA1 (3B11NB, 1:50), GZMB (EPR20129-217, 1:100), H3K27ac (6E7A9, 1:100), H3K4 (C42D8, 1:100), HIF1α (ESEE122, 1:200), HLA-DR (TAL1B5, 1:200), KI67 (EPR3610, 1:200), NFκB (D14E12, 1:100), NLGN1 (1C9.1, 1:100), NLGN4X (S98-7, 1:100), NRXN1 (ABN161-I, 1:100), TUBB3 (TUJ1, 1:100), TOMM20 (ST1705, 1:100), TRYP (AA1, 1:1000), TH (bs-0840R, 1:50), VIM (V9, 1:1000). Alexa Fluor Plus 555 goat anti-mouse (Thermo Scientific, cat no: A32727, 1:100 dilution) or Alexa Fluor Plus 647 goat anti-rabbit (Thermo Scientific, cat no: A32733, 1:200 dilution) secondary antibodies were used to detect mouse or rabbit primary antibodies.

All the primary, secondary antibodies, and DAPI were diluted in Multistaining Buffer (BU06, Lunaphore Technologies). For all staining cycles, the incubation time of primary antibody mixtures was 4 min, and the incubation time of secondary antibodies and DAPI mixtures was 2 min, both at 37 °C. Then, slides in Imaging Buffer (BU09, Lunaphore Technologies) were imaged at 20X magnification. For each imaging cycle, exposure times of 25, 400, and 250 ms were used for DAPI, TRITC, and Cy5 channels, respectively. Samples were eluted for 2 min with elution Buffer (BU07-L, Lunaphore Technologies) at 37 °C and quenched using quenching buffer (BU08-L, Lunaphore Technologies) at 37 °C for 30 s before the next cycle.

### Data Preprocessing

IF data were log-transformed and clipped at the 5th and 99.9th percentiles for CRC data and 0.1th and 99.9th percentiles for prostate data before being rescaled to [0,255]. Whole cell segmentation was performed using Mesmer^51^ using the DAPI channel as the nuclear marker and a maximum projection of the CD45 and PanCK channels as the membrane marker. Image patches were then cropped around each segmented cell with a size of 32×32 pixels at a resolution of 0.65 microns per pixel. Image registration was performed to align H&E and IF images. First, the H&E image was deconvolved into separate Hematoxylin and Eosin channels. Then, the Hematoxylin channel was registered to the DAPI channel from the IF image by performing an affine transformation, calculated by identifying keypoint matches between the images using the OpenCV implementation of Scale Invariant Feature Transform (SIFT) feature extraction^52^.

For the prostate cohort, preprocessing followed the same pipeline as the CRC data, including log transformation, percentile clipping, rescaling, cell segmentation, patch extraction, and H&E-IF registration, with minor adjustments. Because H&E was acquired at the end of the protocol, some regions exhibited partial tissue loss and therefore lacked corresponding cells between modalities; QC-based filtering was applied to exclude these tissue-poor regions.

### Tokenizer Training

To tokenize paired H&E and multiplex IF images, we considered two approaches: one single VQGAN for both modalities, and separate VQGANs trained for each modality. For training, we used default settings apart from four downsampling steps, resulting in 4×4 quantized latent representations. To optimize VQGAN tokenizer training for IF-to-H&E image synthesis, we systematically evaluated codebook size and architectural configurations (Supp. Table 1).

We first trained IF-only tokenizers with codebook sizes of 256 and 1024. This is done by treating each individual marker channel as an independent single-channel image, resulting in a marker-agnostic tokenization approach. Both models achieved high marker-level fidelity, with Spearman correlations of 0.98 and 0.95 and structural similarity index (SSIM) values of 0.77 and 0.82, respectively.

We next trained joint IF-H&E tokenizers using a shared codebook (IF-HE-256 and IF-HE-1024). This is done by deconvolving the H&E into separate Hematoxylin and Eosin channels and treating these as two additional IF marker channels. These models achieved comparable performance to IF-only tokenizers, with Spearman correlations of 0.98 and SSIM values ranging from 0.78-0.80, indicating that joint tokenization did not compromise reconstruction accuracy.

To assess the effect of modality-specific capacity, we further explored configurations where IF and H&E tokenizers were trained independently with different codebook sizes. In this approach we keep H&E images in their original RGB format. The IF-HE-256,256 configuration (separate IF and HE tokenizers, each with 256 codebook size) achieved the best overall performance, combining high marker-level correlation (Spearman: 0.96) with substantially improved structural fidelity (SSIM: 0.87). Reducing IF capacity while maintaining H&E capacity (IF-HE-128,256) resulted in a moderate performance decrease (Spearman: 0.94, SSIM: 0.84). In contrast, the low-capacity configuration (IF-HE-128,128) showed marked degradation in both correlation (Spearman: 0.52) and structural similarity (SSIM: 0.78), indicating insufficient representational capacity.

Based on these results, we selected separate IF & HE tokenizers with 256 codebook size (IF-HE-256,256) for all downstream analyses. This configuration consistently preserved high marker-level correlations (>0.95 for CD31, CD45, CD68, and FOXP3) while achieving superior structural similarity, supporting effective learning of discriminative and spatially coherent representations across diverse marker expression patterns.

### Masked Language Model Training

H&E and IF images were tokenized using their respective pretrained VQGAN by flattening the 4×4-dimensional latent representations for each IF channel and H&E image and concatenating each 16-token sequence together. A joint vocabulary of size 512 was created by concatenating the H&E vocabulary to the IF vocabulary. We utilized the RoBERTa architecture from HuggingFace with 24 layers, 16 heads, and 1024 latent dimensions. Three learnable embedding layers were used: token embeddings to embed the token ID, position embeddings to encode the spatial position of the token in each image, and marker embeddings to encode the specific marker channel or H&E image. The learned marker relationships are depicted in Supp. Fig. 6. During training, a masking ratio is randomly sampled from (0,1) to determine the number of markers to be masked, and for each masked marker, all 16 tokens corresponding to that marker channel are masked. For CRC data, we utilized a training set of 39 patient samples, using a batch size of 64 and evaluate results on 9 held-out samples (CRC01, CRC04, CRC07, CRC10, CRC13, CRC19, CRC23, CRC32, CRC36, CRC40). For prostate cancer data, we utilized a training set of 34 samples and a batch size of 32. We conducted training using eight Nvidia A100 GPUs.

### Iterative Panel Selection Algorithm

We employed a greedy selection algorithm to determine an ordering of biomarkers based on their informativeness for predicting the remaining markers. Specifically, given a selected set of test images, the algorithms identifies an ordered sequence of markers *M* such that for *N* markers, the input set of markers {*m*_1_,…,*m*_k_} is the best set of *k* markers that produce the remaining markers {*m_N_*,…,*m*_k_}. We call this procedure iterative panel selection (IPS)^34^.

In our implementation, we fix *m*_1_ to be H&E and iteratively expand the marker set from *m*_2_ to *m_k_* using a “most informative marker first” heuristic. At each iteration, we chose the marker whose inclusion maximizes prediction performance for the remaining markers. This greedy approach yields an ordered list in which earlier markers contribute the greatest marginal gain in overall prediction accuracy.

We additionally considered an alternative ordering based on a complementary “hardest to predict marker first” heuristic. In this case, we determined the ordering in reverse by first identifying the marker that is easiest to predict from the remaining set, corresponding to the final element in the ordering, i.e., finding *m_N_* first, which corresponds to the marker *m* that is easiest to predict from the remaining set of markers *M\m*. We refer to this strategy as reverse iterative panel selection (rIPS)^29^. Together, IPS and rIPS provide complementary perspectives on marker importance and predictability within the panel.

After model training, we utilized a subset of the training dataset to conduct IPS and rIPS. For CRC, we selected two patient samples (CRC02, and CRC03) and clustered cells using K-means clustering. We selected 15 clusters and sample 1000 cells from each cluster to produce a set of 15,000 cells for panel selection.

For the prostate dataset, we sought to evaluate the impact of using different sets of cells to perform panel selection on the final marker ordering. To generate representative subsets for the prostate dataset, we pooled cells from all patients spanning the full range of Gleason grades and performed k-means clustering with a fixed number of clusters (K=15). Using this shared clustering, we constructed three candidate cell orderings by sampling from the same 15 clusters at progressively increasing density (Supp. Fig. 7).

For ordering 1, we sampled 170 cells per cluster (15×170). For ordering 2, we increased the sampling density to 300 cells per cluster (15×300=4,500 cells) to capture greater within-cluster variability while preserving the same cluster structure. For ordering 3, we applied proportional sampling, in which the number of cells selected from each cluster was proportional to its cluster size (still with K=15). This produced a larger representative set of 9,000 cells that preserves original cluster abundance and more accurately reflects the balance between rare and dominant cellular phenotypes across patients and Gleason grades.

### UNI/KRONOS Benchmark

We benchmarked our H&E-MTI model against two foundation models, UNI2-h and KRONOS, for the prediction of immunofluorescence (IF) marker intensities.

To evaluate H&E-only performance, we compared against UNI2-h, a Vision Transformer pretrained on over 200 million pathology H&E and IHC images from approximately 350,000 whole-slide images (WSIs) using the DINOv2 self-supervised learning framework. For feature extraction, H&E channels were extracted from CRC ORION images and passed through the frozen UNI2-h encoder to generate 1,536-dimensional embeddings. A regression model was trained on these embeddings to predict mean intensities of 14 IF markers, and performance was evaluated using Spearman correlations between predicted and ground-truth marker mean intensities, assessing UNI2-h’s ability to predict IF marker expression from morphology alone.

We also benchmarked against KRONOS^40^, a foundation model specifically designed for spatial proteomics. KRONOS is a Vision Transformer pretrained on over 47 million multiplexed imaging patches spanning 175 protein markers and 8 imaging platforms (CODEX, COMET, IBEX, MxIF), with architectural adaptations to handle multi-channel data, unlike general histopathology models trained solely on H&E images. To evaluate KRONOS’s ability to predict IF marker expression from an optimized input panel, we extracted three IF marker channels from our CRC ORION images and pass them through the frozen KRONOS encoder to generate 1,152-dimensional feature embeddings (3 markers × 384 dimensions). A regression model was trained to predict mean intensities of the remaining 14 IF markers, with performance evaluated using Spearman and Pearson correlations between predicted and ground-truth values across multiple CRC samples.

To test whether combining morphological and partial molecular information improves prediction performance, we developed a combined UNI-KRONOS model that fuses features from both modalities. We extracted 1,536-dimensional embeddings from H&E images using the frozen UNI2-h encoder and 1,152-dimensional embeddings from three IF marker channels (CD8a, CD163, PD-1; 3 marker x 384 dimensions) using the frozen KRONOS encoder. These embeddings were concatenated to form 2,688-dimensional feature vectors capturing complementary morphological and partial immunofluorescence information. A multi-layer regression model with batch normalization and dropout was trained to predict mean intensities of the remaining 14 IF markers.

Finally, we benchmarked these approaches against our proposed miniMTI method. For this benchmark, we provided the model with H&E images augmented by a minimal set of experimentally acquired IF markers selected using our iterative panel optimization strategy. The miniMTI model tokenized H&E and sparse IF inputs and predicted the remaining IF marker intensities using the same evaluation protocol above. Performance was assessed using Spearman and Pearson correlations between predicted and ground-truth mean marker intensities across CRC samples. This benchmark directly evaluated whether sparse molecular anchoring via miniMTI provides improved predictive power compared to H&E-only models, proteomics-native encoders with partial input, or fusion approaches.

### Spatial tissue organization analysis with CellCharter

To evaluate whether miniMTI preserves spatial tissue architecture, we performed CellCharter-based spatial clustering on both ground-truth and miniMTI-imputed marker data from nine colorectal cancer samples in the CRC-ORION cohort (CRC04, CRC07, CRC10, CRC13, CRC19, CRC23, CRC32, CRC36, CRC40). CellCharter (v1.0) integrates molecular and spatial information to identify tissue neighborhoods defined by coordinated cell-type composition and spatial proximity, making it well-suited for evaluating spatial fidelity of imputed marker profiles as well as their cell phenotypes.

For each sample, we constructed two AnnData objects (Scanpy v1.9), one containing experimentally measured marker intensities and the other containing corresponding miniMTI-imputed values. Analyses were performed on 14 markers (DAPI, CD31, CD45, CD68, CD4, FOXP3, CD45RO, CD20, PD-L1, CD3e, E-cadherin, Ki67, PanCK, and αSMA). Marker intensities were z-score normalized with outlier clipping at ±10 standard deviations.

Spatial graphs were constructed using Delaunay triangulation (Squidpy v1.3.0) to connect neighboring cells, followed by removal of edges exceeding the 99th percentile of all edge lengths to eliminate spurious long-range connections at tissue boundaries. For each cell, we performed 3-layer neighborhood aggregation by concatenating the cell’s own marker profile with mean marker profiles of cells at 1, 2, and 3 steps away in the spatial graph. This representation captured local tissue microenvironments by jointly encoding intrinsic cellular states and surrounding neighborhood composition.

Spatially aggregated features were clustered using a Gaussian Mixture Model (n_clusters=8, random_state=42) implemented in PyCave (v3.2.1) on CPU to identify spatial niches. For each sample, cluster labels were derived from ground-truth and imputed data were aligned using the Hungarian algorithm (scipy.optimize.linear_sum_assignment) based on maximum spatial overlap quantified from the confusion matrix. Agreement between ground-truth and imputed spatial domains was quantified using the Adjusted Rand Index (ARI).

To characterize cluster-level molecular consistency, we computed mean z-scored marker expression intensities for each cluster and evaluate cell-level Pearson correlation coefficients between ground-truth and imputed marker expression patterns within corresponding spatial domains.

### Quantifying preservation of dynamic range

To evaluate whether imputation preserves both continuous marker intensity distributions and the fraction of marker-positive cells, we assessed real and imputed data using a shared, real-derived positivity threshold for each marker. For each marker, we fit a two-component Gaussian Mixture Model (GMM) to the distribution of real single-cell intensities, modeling negative and positive populations, and define the positivity threshold at the GMM decision boundary.

This threshold, derived from the real data, was held fixed and applied unchanged to the corresponding imputed intensities to binarize cells into positive and negative classes. Using these binary assignments, we computed true positives (TP), false positives (FP), true negatives (TN), and false negatives (FN), and summarized agreement between real and imputed calls using row-normalized confusion matrices.

In parallel, to assess preservation of dynamic range beyond binary classification, we overlayed real and imputed single-cell intensity histograms for each marker, with the positivity threshold indicated. This comparison enabled visual evaluation of whether imputed signals recapitulate the full intensity spectrum and distributional shape of the corresponding real measurements, rather than merely matching binary positivity rates.

### Quantifying preservation of disease progression-marker signatures

For markers selected a priori for their known association with prostate cancer progression (VIM, AMACR, αSMA, and Tryptase), we assessed whether virtual staining preserved disease-linked marker relationships by comparing positivity trends across Gleason grade groups in real and imputed data. Per-cell positive and negative labels were obtained using the GMM-thresholding procedure described above.

For each patient and Gleason category (no cancer, Gleason 3+3, 3+4, and 4+3), we computed the fraction of marker-positive cells. These per-sample positive rates were summarized using grouped boxplots stratified by Gleason grade and input condition (Real staining versus H&E-based prediction). Agreement was evaluated by examining whether the imputed data recapitulate the same grade-dependent shifts in marker positivity observed in the corresponding real measurements.

For each marker, we further quantified concordance using two complementary metrics. First, we computed Pearson’s correlation coefficient (r) between real and predicted per-sample positive fractions across patients, providing a direct measure of sample-level agreement. Second, to explicitly assess preservation of Gleason-associated progression patterns, we performed β-distribution fitting by using the mean positivity within each Gleason group as summary points and estimating a single pair of β parameters for the real data and for the imputed data. The resulting parameter estimates were then compared between conditions.

For a proportion *x* ∈ (0, 1) with shape parameters *a* > 0 and *b* > 0, the β-distribution is defined as:

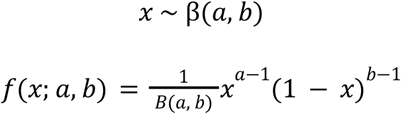

where the normalization constant *B*(*a, b*) is the Beta function:

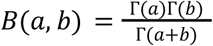

where Γ(.) denotes the Gamma function.

This analysis specifically evaluates preservation of disease progression-associated marker signatures (Supp. Fig. 5), rather than overall intensity similarity alone, providing a clinically relevant assessment of whether virtual staining maintains biologically meaningful trends across disease states.

### Spatial Auto-correlation Analysis (Moran’s I)

We quantified global spatial autocorrelation using Moran’s I for each protein marker in both the original multiplex tissue imaging (MTI) data and the corresponding imputed predictions. Moran’s I is a statistic that measures the degree to which similar expression values are spatially clustered, with higher values indicating stronger global spatial organization.

For each tissue sample, Moran’s I was computed on a per-marker basis using cell-level spatial coordinates. To compare spatial autocorrelation between original and imputed data while accounting for repeated measurements across samples and markers, we fit linear mixed-effect models with Moran’s I as the response variable.

## Data availability

Due to the large size of the dataset and ongoing analysis, the 40-plex MTI and matched H&E data generated in this study from 63 prostate cancer patients are available from the corresponding author upon reasonable request.

## Code availability

All code used for data processing, model training, and evaluation is available at https://github.com/ohsu-cedar-comp-hub/cycif-panel-reduction. Model weights are available upon request.

## Supplementary Information

**Supplementary Figure 1** Additional visualizations of generated images. Top 6 images from the CRC dataset and bottom 6 images from the prostate dataset.

**Supplementary Figure 2 a**, Marker-wise comparison of prediction performance (Pearson correlation) across models, including foundational model ensembles (UNI, KRONOS, and UNI+KRONOS) and miniMTI configurations using IF-only inputs, and IF plus H&E inputs. **b**, Sample-wise comparison of the same models across nine held-out colorectal cancer patient samples.

**Supplementary Figure 3.** CellCharter spatial clustering maps from Ground Truth MTI and miniMTI-predicted intensities using H&E+3 markers for all CRC validation samples

**Supplementary Figure 4**. Full marker-wise intensity distributions (real vs imputed) and real-derived positivity gating across input panel sizes (H&E-only, H&E+3, H&E+6, H&E+9) in the prostate cohort.

**Supplementary Figure 5**. β-distribution fitting of Gleason-stratified marker-positive fractions for progression-associated markers (VIM, AMACR, αSMA, Tryptase), comparing Real vs H&E-only vs H&E+3.

**Supplementary Figure 6** a. tSNE plot of marker embeddings from the prostate model. Colors represent clusters assigned via k-means clustering. b. Heatmap depicting the cosine similarity between marker embeddings from the prostate model. c. tSNE of marker embeddings from the CRC model. d. Heatmap depicting the cosine similarity between markers in the CRC model.

**Supplementary Figure 7**. Stability of the prostate marker ordering across three IPS runs using three different subsampled cell sets (orderings 1, 2, 3).

**Supplementary Table 1**. Hyperparameter optimization for VQGAN tokenizer training across architectural configurations

