## Supplementary for "miniMTI: minimal multiplex tissue imaging enhances biomarker expression prediction from histology"

### Supplementary Figure 1

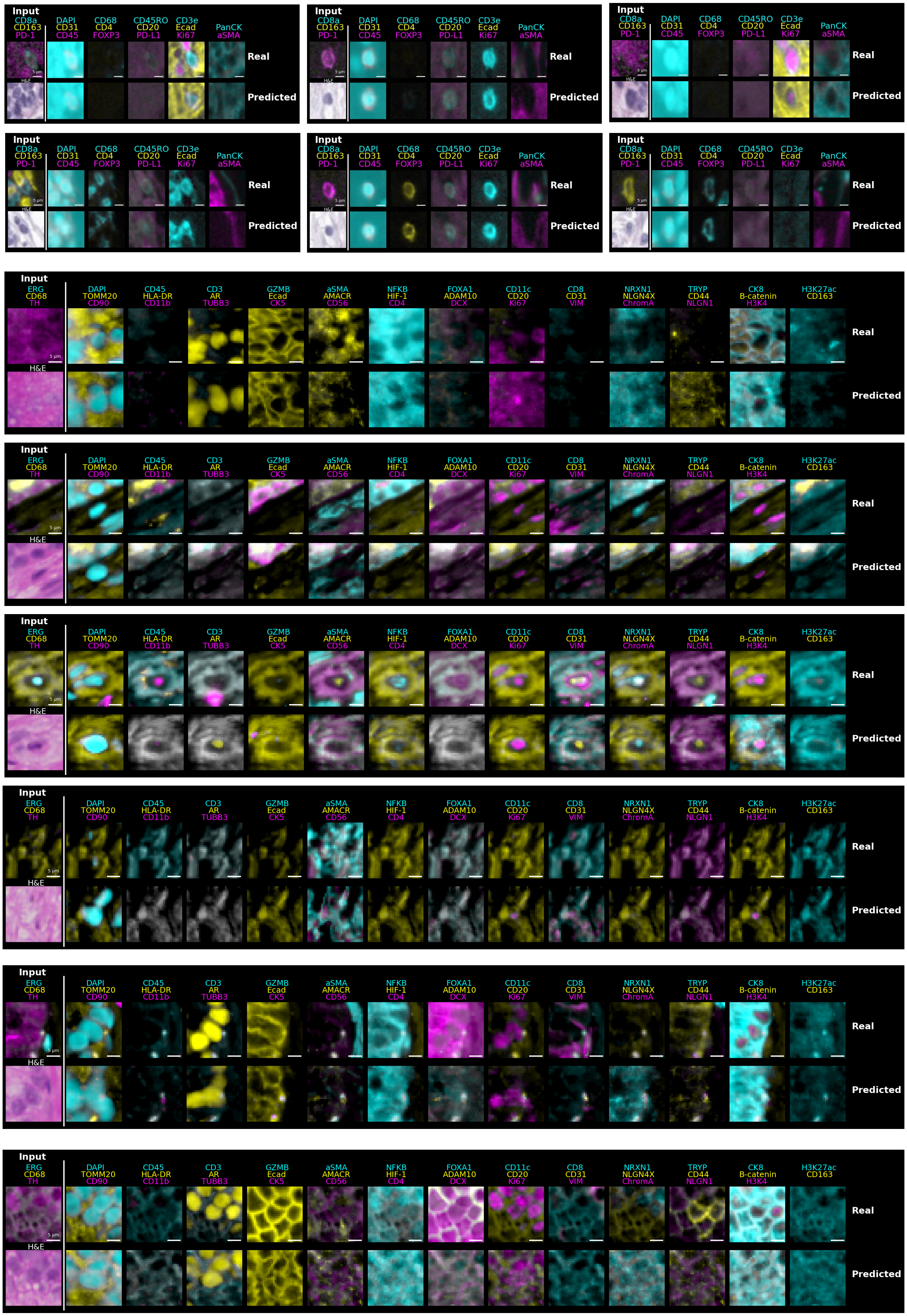

#### Supplementary Figure 2

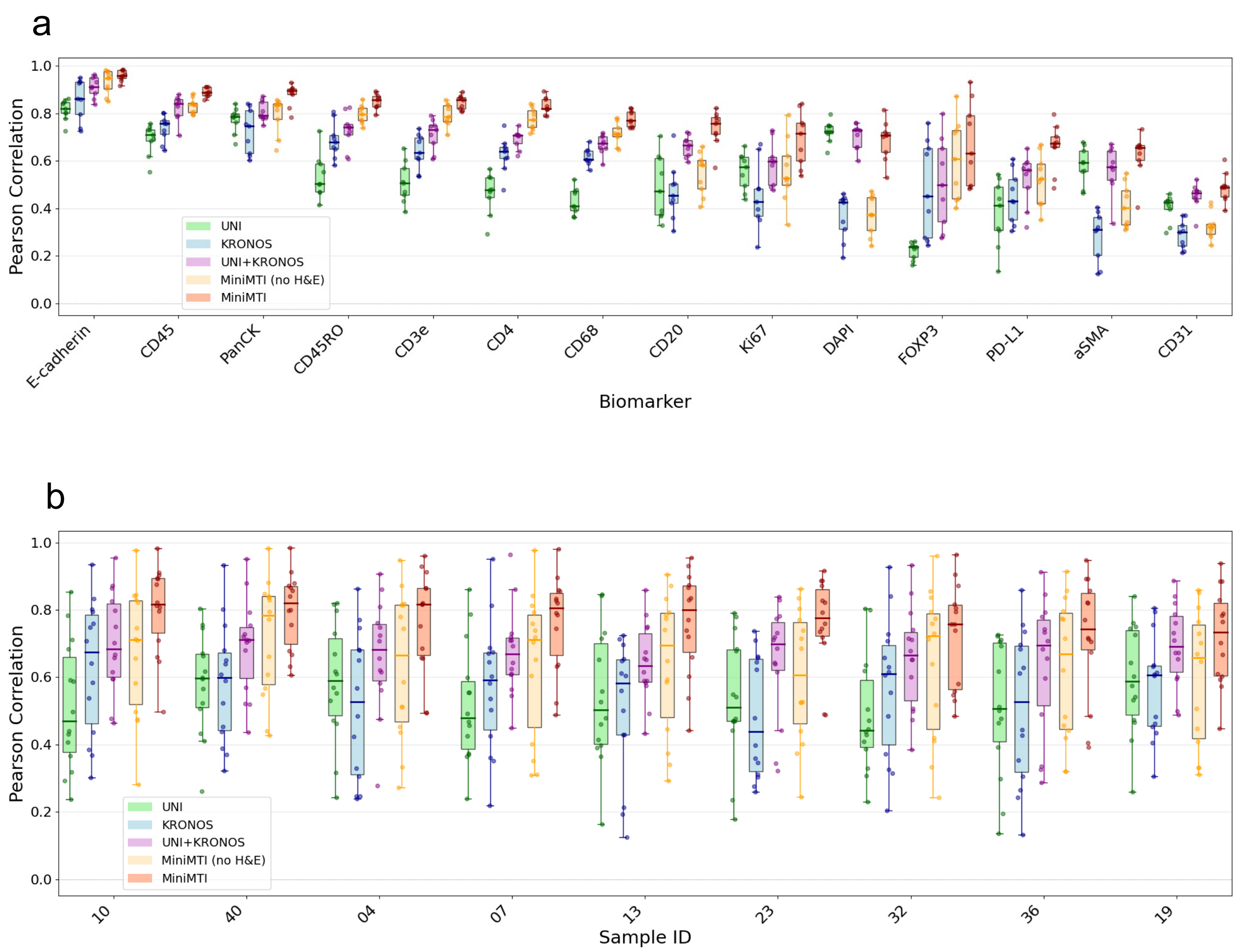

### Supplementary Figure 3

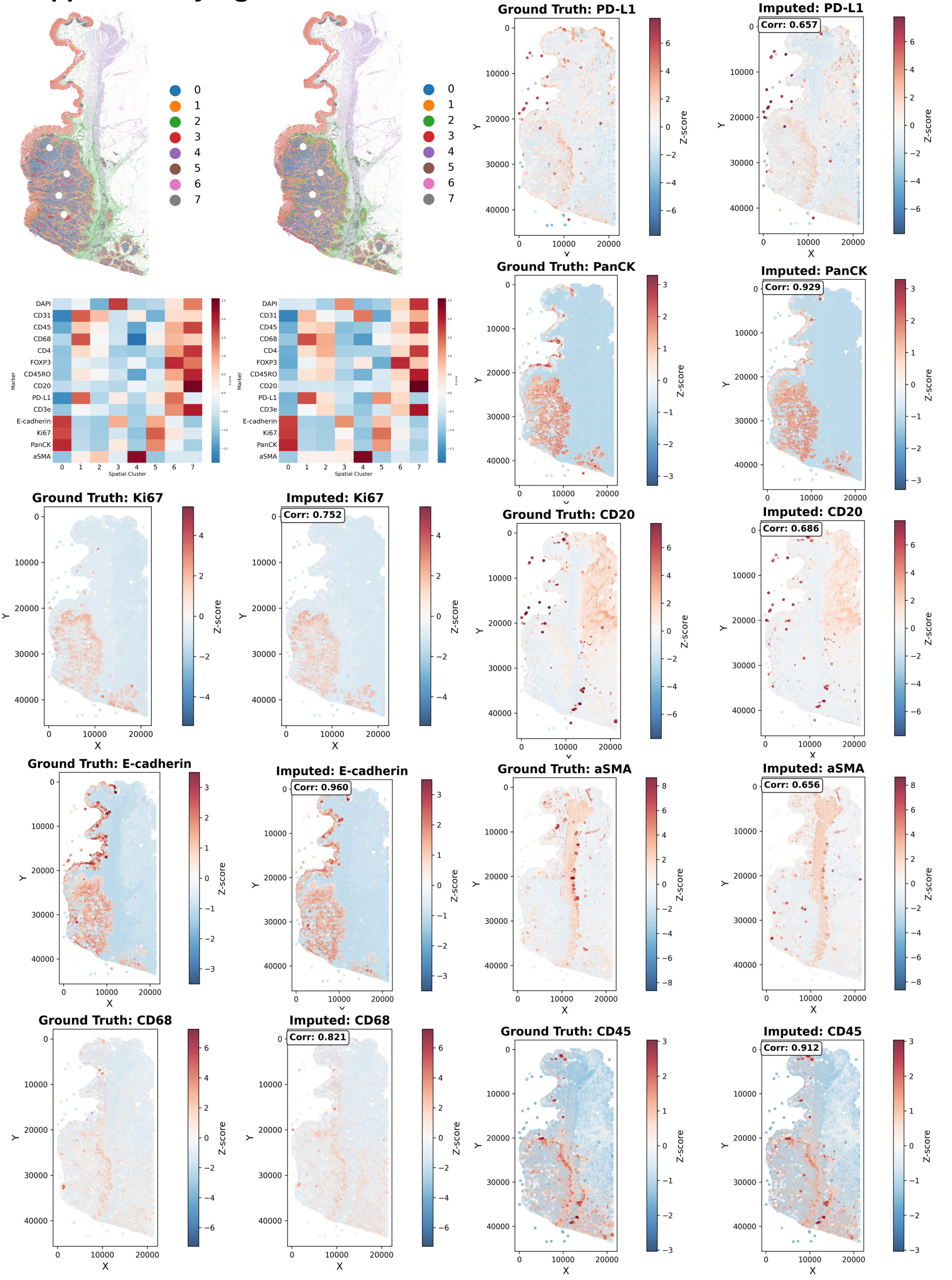

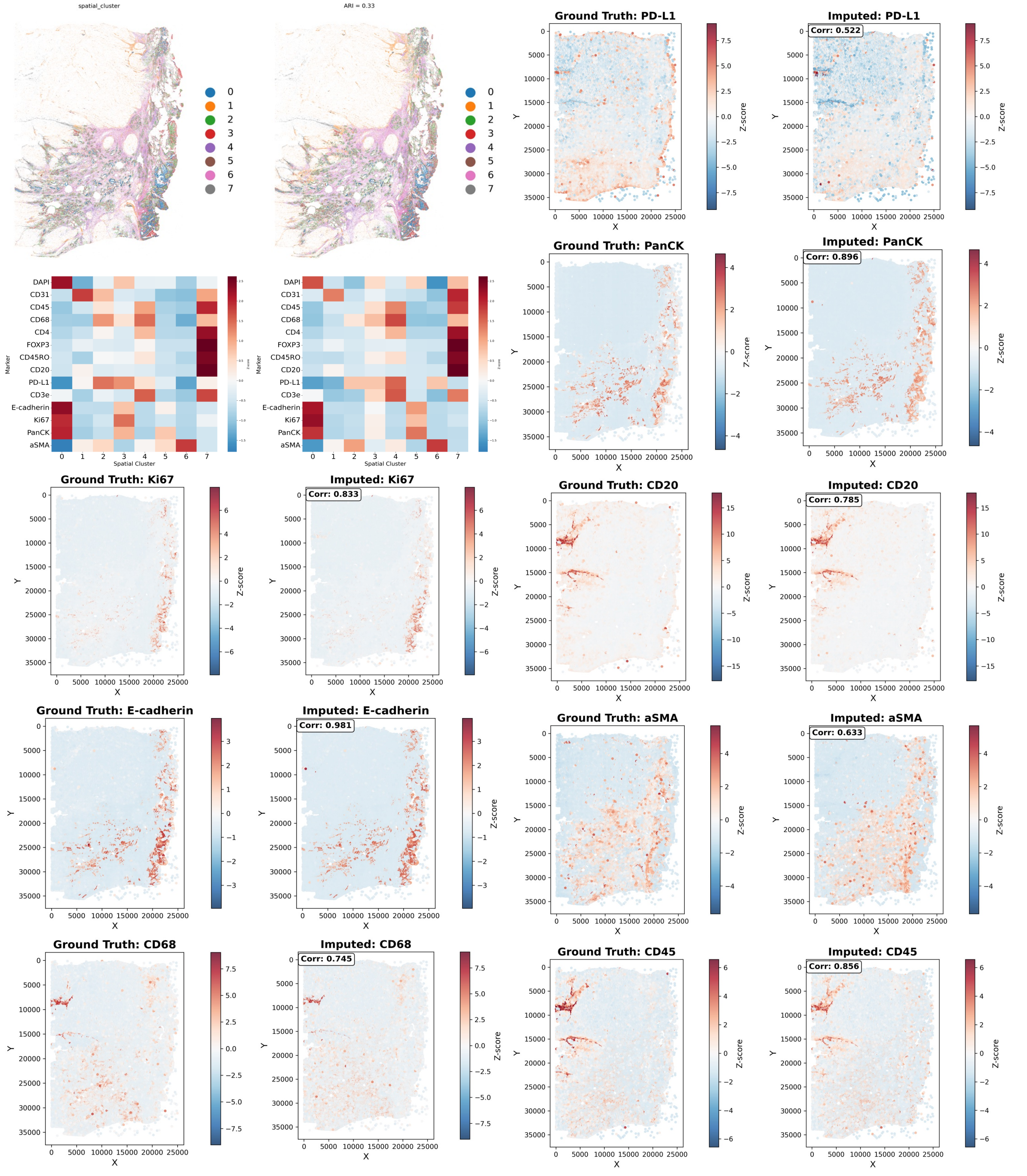

spatial_cluster
ARI = 0.33
Spatial Cluster
Marker
DAPI
CD31
CD45
CD68
CD4
FOXP3
CD45RO
CD20
PD-L1
CD3e
E-cadherin
Ki67
PanCK
aSMA
Ground Truth: PD-L1
Imputed: PD-L1
Corr: 0.522
Ground Truth: PanCK
Imputed: PanCK
Corr: 0.896
Ground Truth: Ki67
Imputed: Ki67
Corr: 0.833
Ground Truth: CD20
Imputed: CD20
Corr: 0.785
Ground Truth: E-cadherin
Imputed: E-cadherin
Corr: 0.981
Ground Truth: aSMA
Imputed: aSMA
Corr: 0.633
Ground Truth: CD68
Imputed: CD68
Corr: 0.745
Ground Truth: CD45
Imputed: CD45
Corr: 0.856
X
Y
Z-score

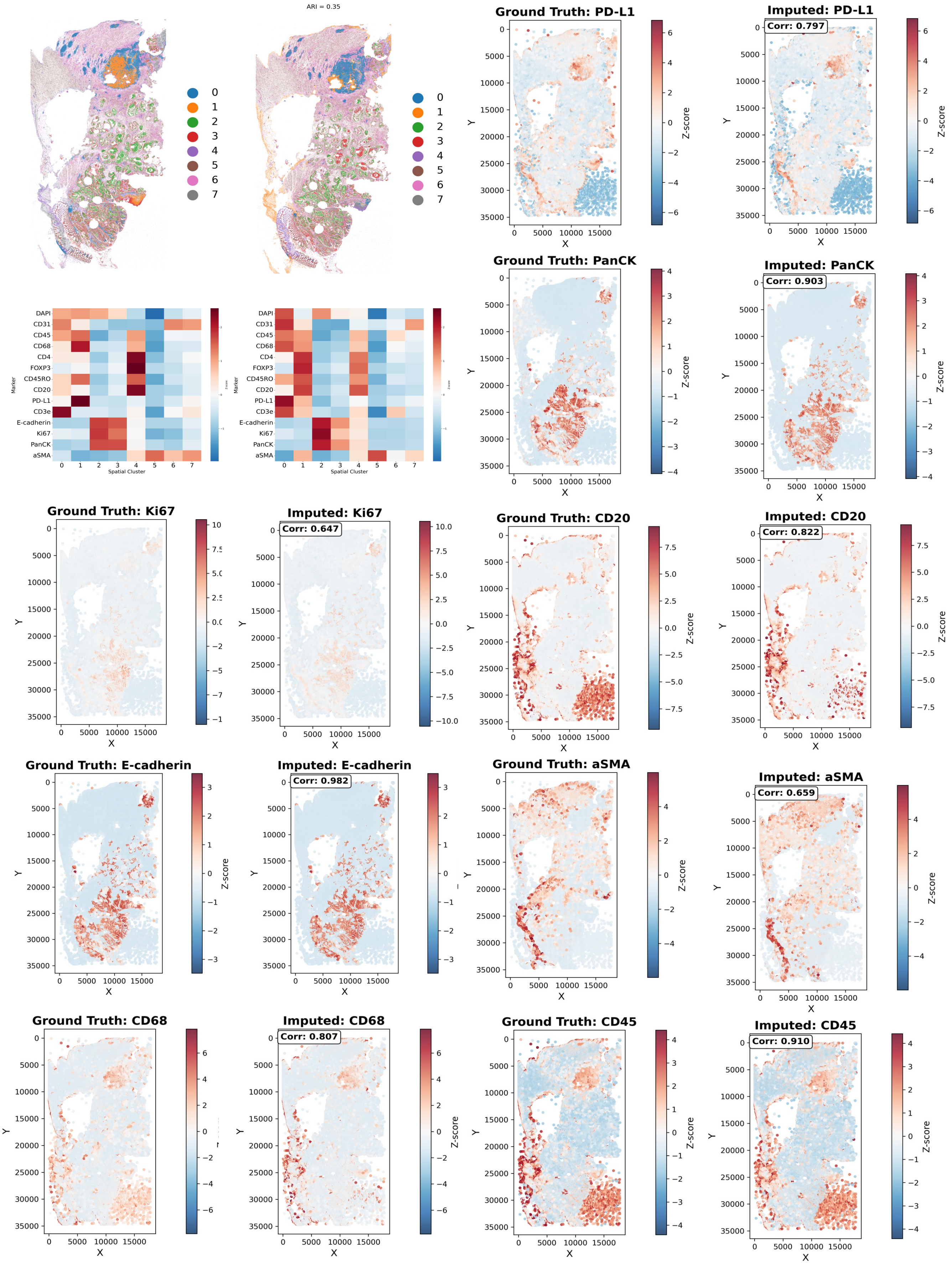
ARI = 0.35
0
1
2
3
4
5
6
7
DAPI
CD31
CD45
CD68
CD4
FOXP3
CD45RO
CD20
PD-L1
CD3e
E-cadherin
Ki67
PanCK
aSMA
Marker
Spatial Cluster
Ground Truth: PD-L1
Imputed: PD-L1
Corr: 0.797
Ground Truth: PanCK
Imputed: PanCK
Corr: 0.903
Ground Truth: Ki67
Imputed: Ki67
Corr: 0.647
Ground Truth: CD20
Imputed: CD20
Corr: 0.822
Ground Truth: E-cadherin
Imputed: E-cadherin
Corr: 0.982
Ground Truth: aSMA
Imputed: aSMA
Corr: 0.659
Ground Truth: CD68
Imputed: CD68
Corr: 0.807
Ground Truth: CD45
Imputed: CD45
Corr: 0.910
X
Y
Z-score

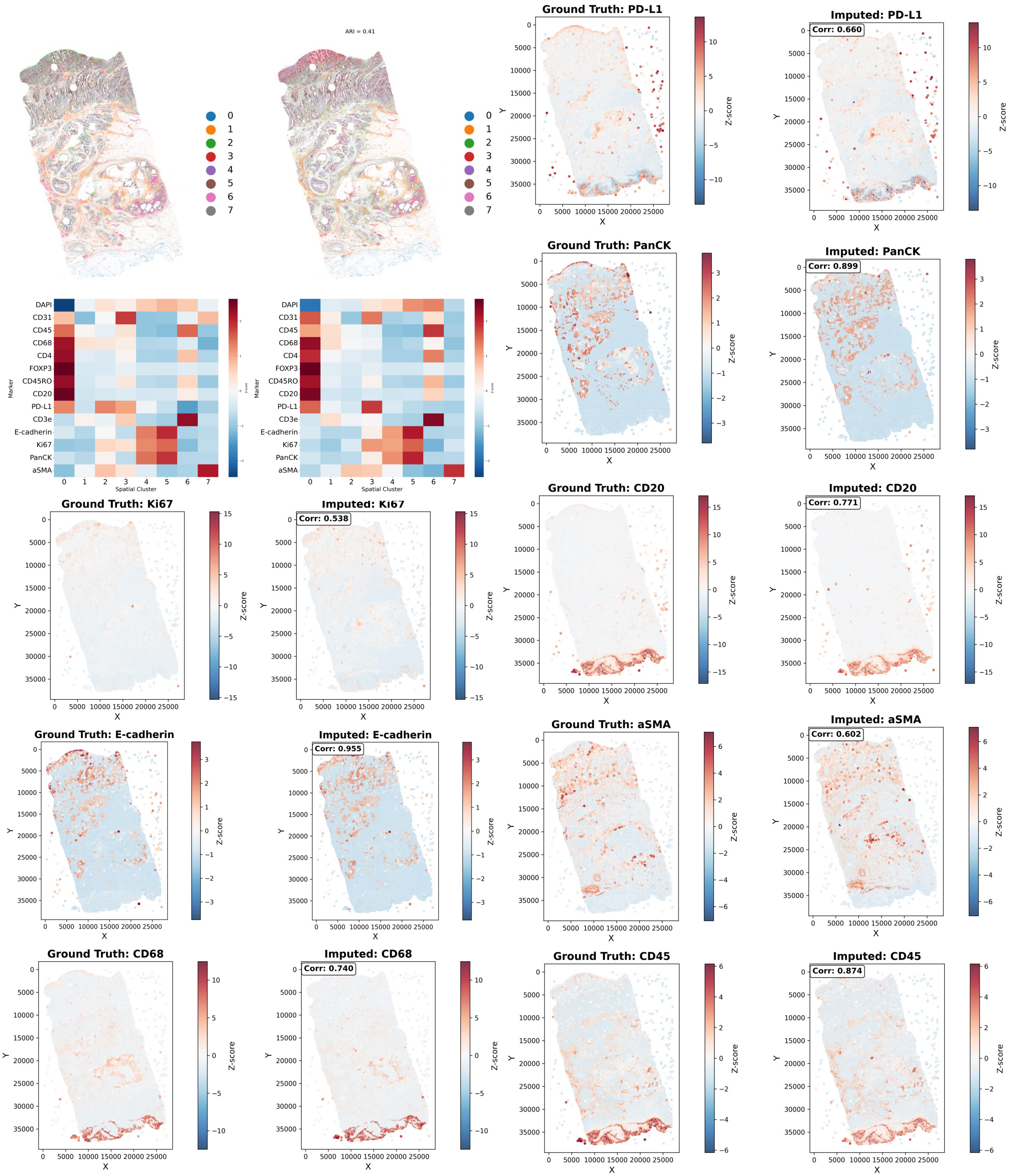

ARI = 0.41
0
1
2
3
4
5
6
7
Marker
DAPI
CD31
CD45
CD68
CD4
FOXP3
CD45RO
CD20
PD-L1
CD3e
E-cadherin
Ki67
PanCK
aSMA
Spatial Cluster
Ground Truth: PD-L1
Imputed: PD-L1
Corr: 0.660
Ground Truth: PanCK
Imputed: PanCK
Corr: 0.899
Ground Truth: Ki67
Imputed: Ki67
Corr: 0.538
Ground Truth: CD20
Imputed: CD20
Corr: 0.771
Ground Truth: E-cadherin
Imputed: E-cadherin
Corr: 0.955
Ground Truth: aSMA
Imputed: aSMA
Corr: 0.602
Ground Truth: CD68
Imputed: CD68
Corr: 0.740
Ground Truth: CD45
Imputed: CD45
Corr: 0.874
X
Y
Z-score

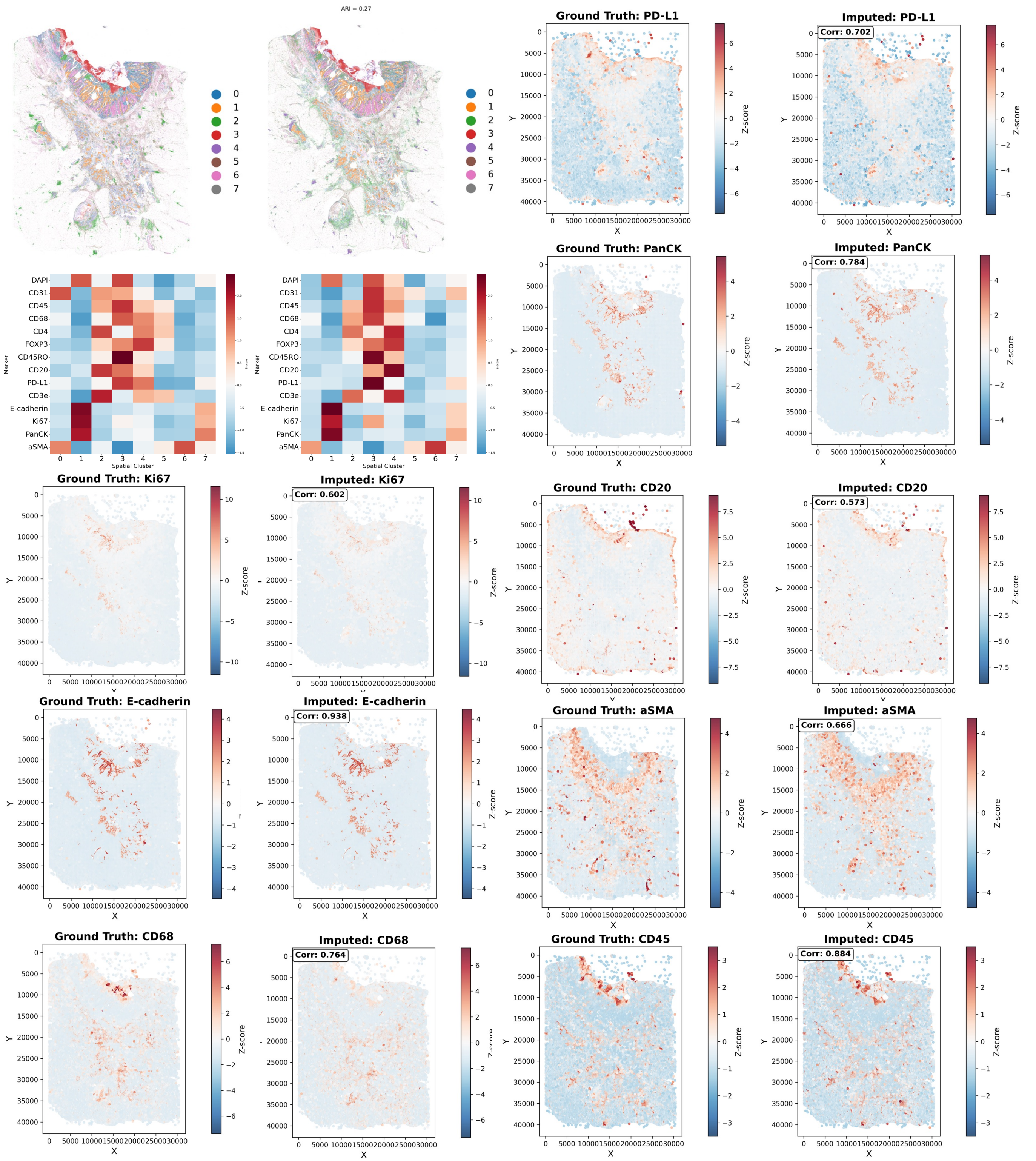
ARI = 0.27
Marker
Spatial Cluster
DAPI
CD31
CD45
CD68
CD4
FOXP3
CD45RO
CD20
PD-L1
CD3e
E-cadherin
Ki67
PanCK
aSMA
Ground Truth: PD-L1
Imputed: PD-L1
Corr: 0.702
Ground Truth: PanCK
Imputed: PanCK
Corr: 0.784
Ground Truth: Ki67
Imputed: Ki67
Corr: 0.602
Ground Truth: CD20
Imputed: CD20
Corr: 0.573
Ground Truth: E-cadherin
Imputed: E-cadherin
Corr: 0.938
Ground Truth: aSMA
Imputed: aSMA
Corr: 0.666
Ground Truth: CD68
Imputed: CD68
Corr: 0.764
Ground Truth: CD45
Imputed: CD45
Corr: 0.884
X
Y
Z-score

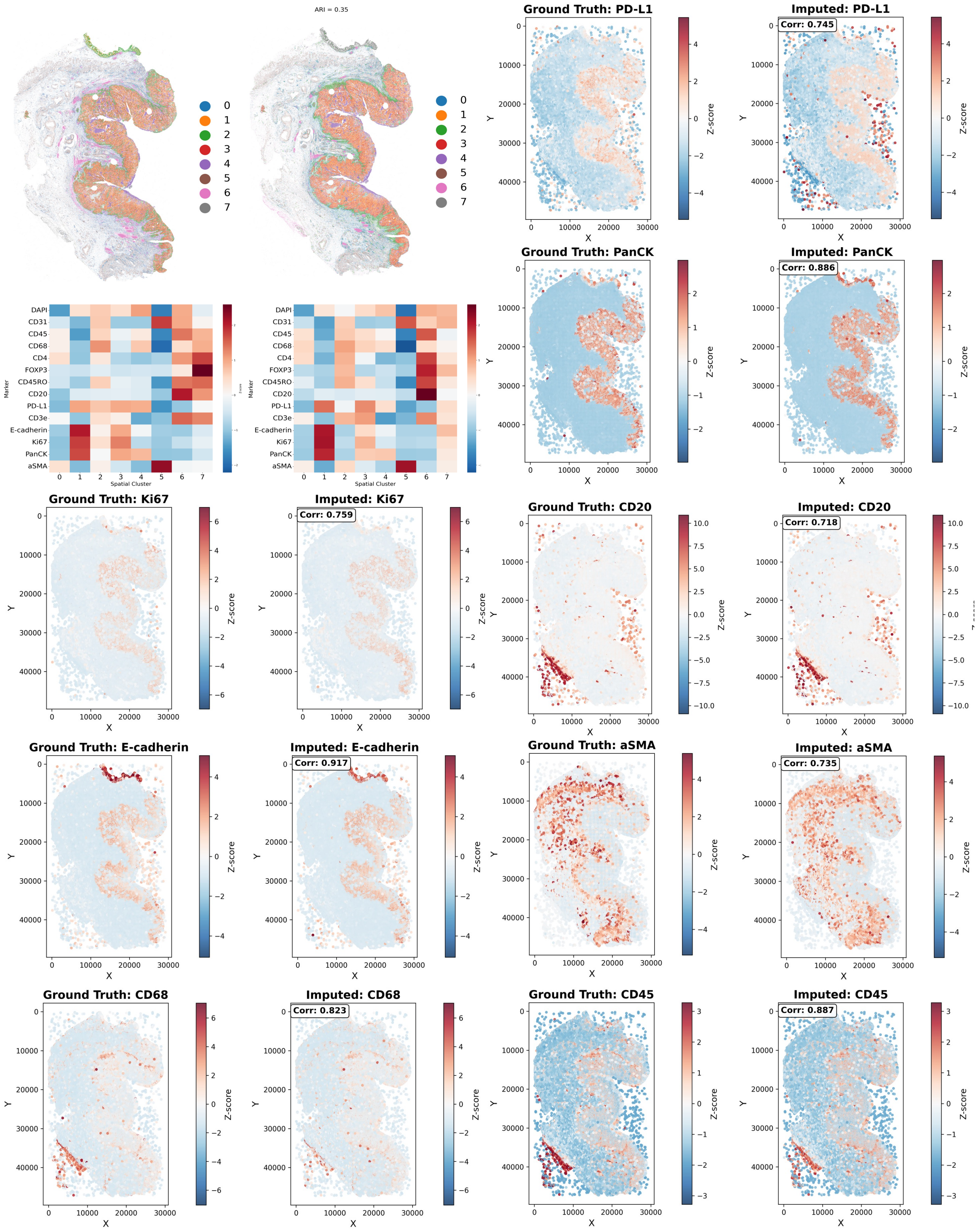

ARI = 0.35
0
1
2
3
4
5
6
7
DAPI
CD31
CD45
CD68
CD4
FOXP3
CD45RO
CD20
PD-L1
CD3e
E-cadherin
Ki67
PanCK
aSMA
Marker
Spatial Cluster
Ground Truth: PD-L1
Imputed: PD-L1
Corr: 0.745
Ground Truth: PanCK
Imputed: PanCK
Corr: 0.886
Ground Truth: Ki67
Imputed: Ki67
Corr: 0.759
Ground Truth: CD20
Imputed: CD20
Corr: 0.718
Ground Truth: E-cadherin
Imputed: E-cadherin
Corr: 0.917
Ground Truth: aSMA
Imputed: aSMA
Corr: 0.735
Ground Truth: CD68
Imputed: CD68
Corr: 0.823
Ground Truth: CD45
Imputed: CD45
Corr: 0.887
X
Y
Z-score

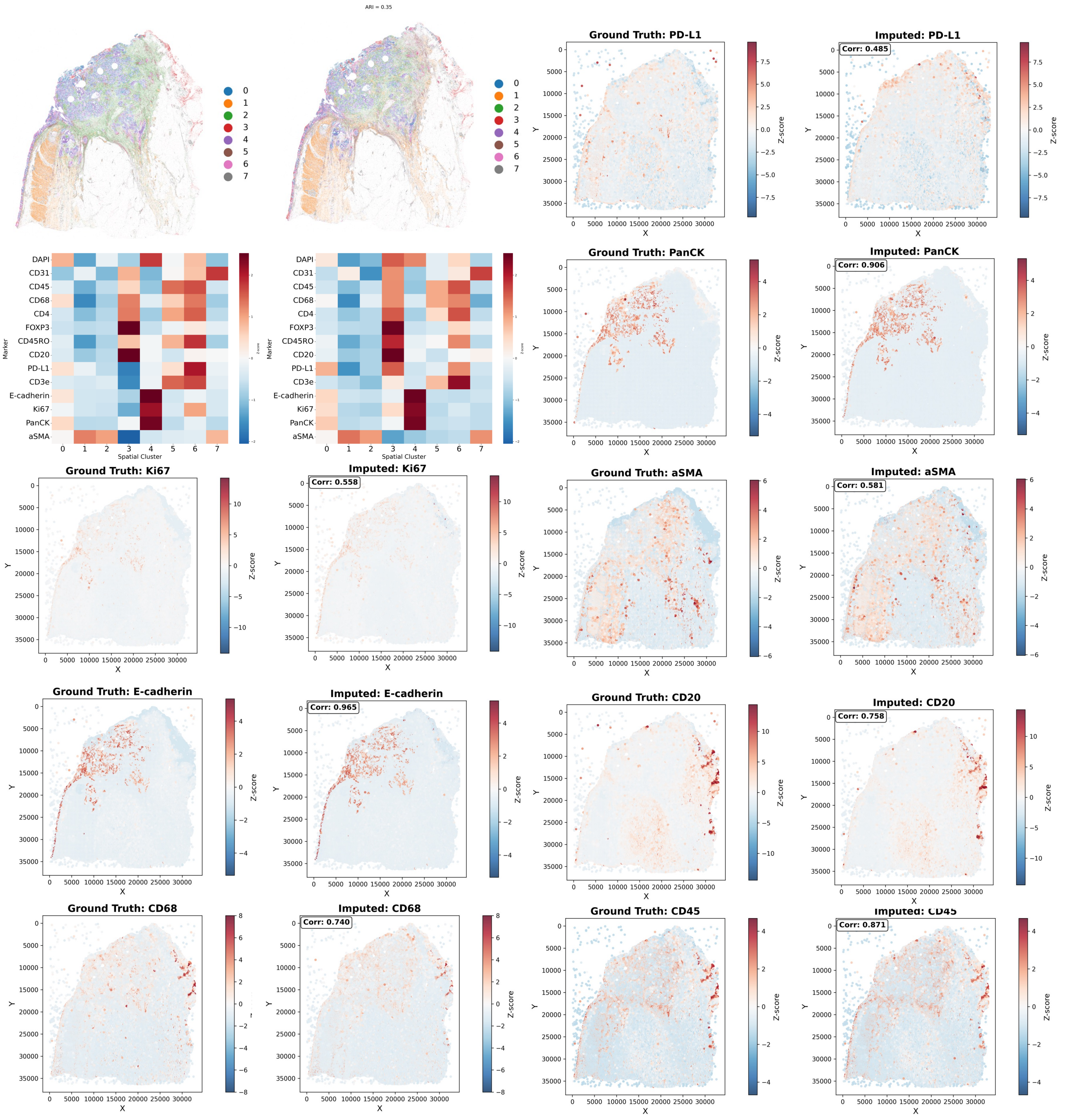
ARI = 0.35
0
1
2
3
4
5
6
7
DAPI
CD31
CD45
CD68
CD4
FOXP3
CD45RO
CD20
PD-L1
CD3e
E-cadherin
Ki67
PanCK
aSMA
Marker
Spatial Cluster
Ground Truth: PD-L1
Imputed: PD-L1
Corr: 0.485
Ground Truth: PanCK
Imputed: PanCK
Corr: 0.906
Ground Truth: Ki67
Imputed: Ki67
Corr: 0.558
Ground Truth: aSMA
Imputed: aSMA
Corr: 0.581
Ground Truth: E-cadherin
Imputed: E-cadherin
Corr: 0.965
Ground Truth: CD20
Imputed: CD20
Corr: 0.758
Ground Truth: CD68
Imputed: CD68
Corr: 0.740
Ground Truth: CD45
Imputed: CD45
Corr: 0.871
X
Y
Z-score

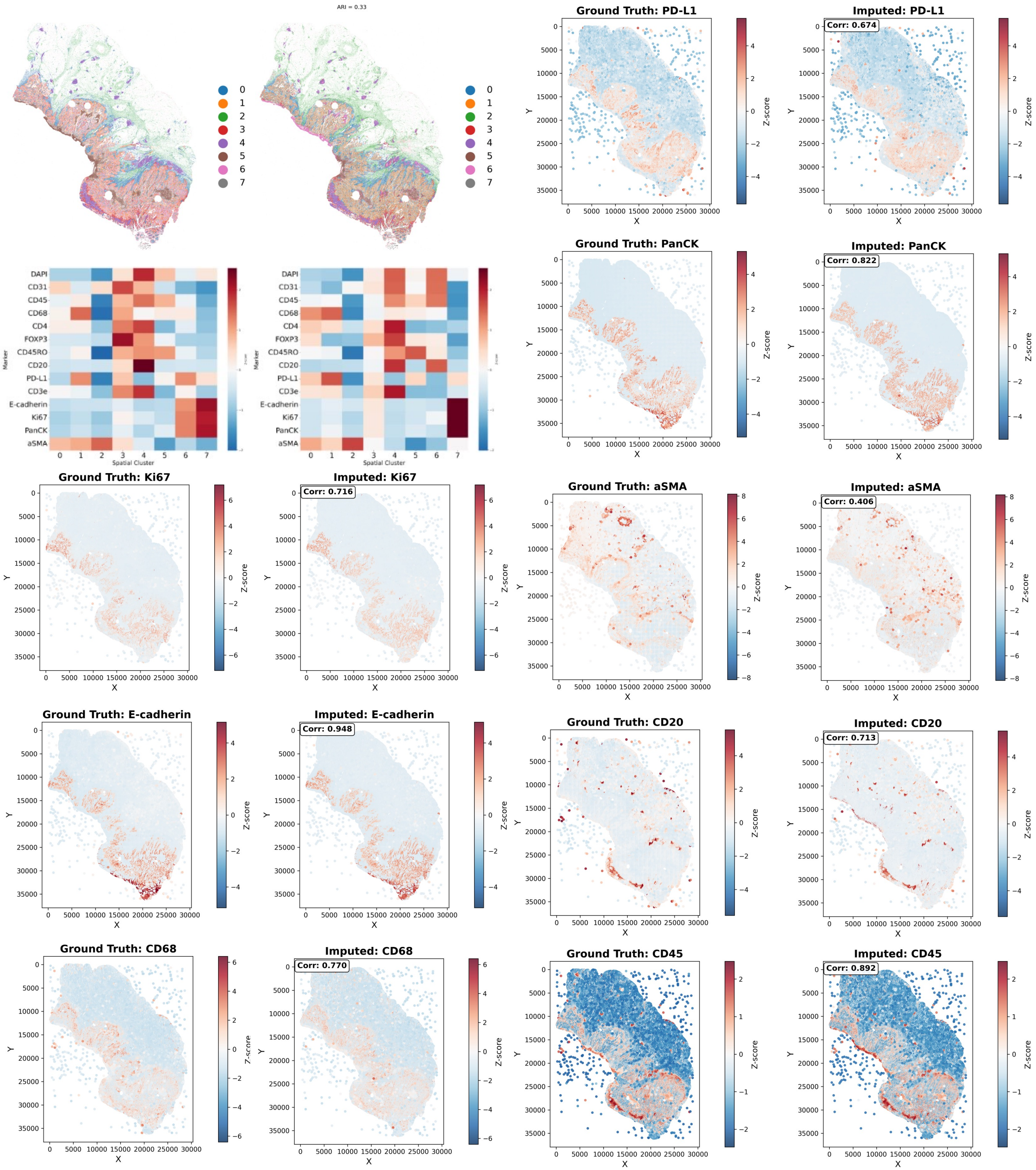
ARI = 0.33
0
1
2
3
4
5
6
7
DAPI
CD31
CD45
CD68
CD4
FOXP3
CD45RO
CD20
PD-L1
CD3e
E-cadherin
Ki67
PanCK
aSMA
Marker
Spatial Cluster
Ground Truth: PD-L1
Imputed: PD-L1
Corr: 0.674
Ground Truth: PanCK
Imputed: PanCK
Corr: 0.822
Ground Truth: Ki67
Imputed: Ki67
Corr: 0.716
Ground Truth: aSMA
Imputed: aSMA
Corr: 0.406
Ground Truth: E-cadherin
Imputed: E-cadherin
Corr: 0.948
Ground Truth: CD20
Imputed: CD20
Corr: 0.713
Ground Truth: CD68
Imputed: CD68
Corr: 0.770
Ground Truth: CD45
Imputed: CD45
Corr: 0.892
X
Y
Z-score

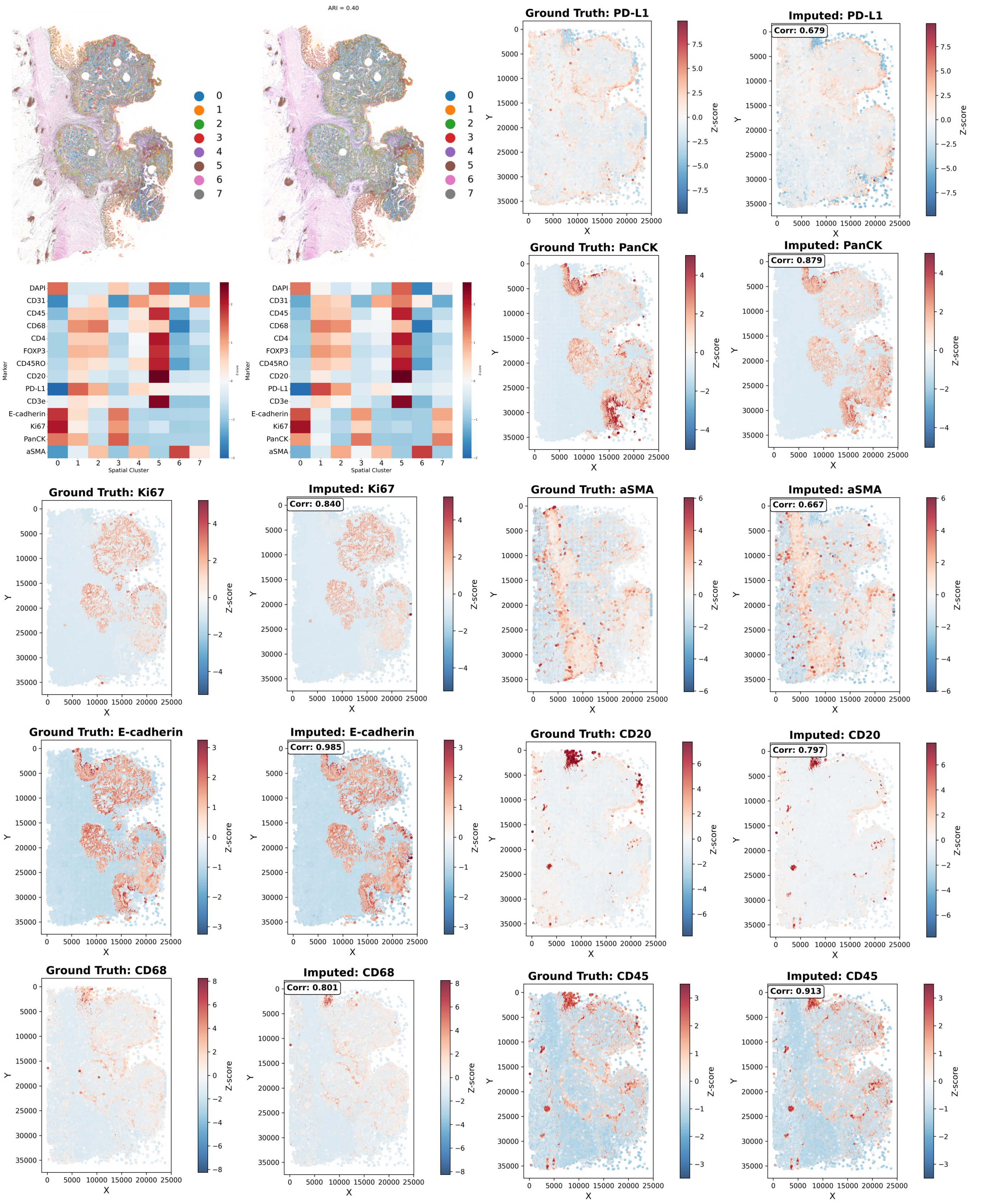
ARI = 0.40
0 1 2 3 4 5 6 7
DAPI
CD31
CD45
CD68
CD4
FOXP3
CD45RO
CD20
PD-L1
CD3e
E-cadherin
Ki67
PanCK
aSMA
Marker
Spatial Cluster
Ground Truth: PD-L1
Imputed: PD-L1
Corr: 0.679
Ground Truth: PanCK
Imputed: PanCK
Corr: 0.879
Ground Truth: Ki67
Imputed: Ki67
Corr: 0.840
Ground Truth: aSMA
Imputed: aSMA
Corr: 0.667
Ground Truth: E-cadherin
Imputed: E-cadherin
Corr: 0.985
Ground Truth: CD20
Imputed: CD20
Corr: 0.797
Ground Truth: CD68
Imputed: CD68
Corr: 0.801
Ground Truth: CD45
Imputed: CD45
Corr: 0.913
X
Y
Z-score

### Supplementary Figure 4

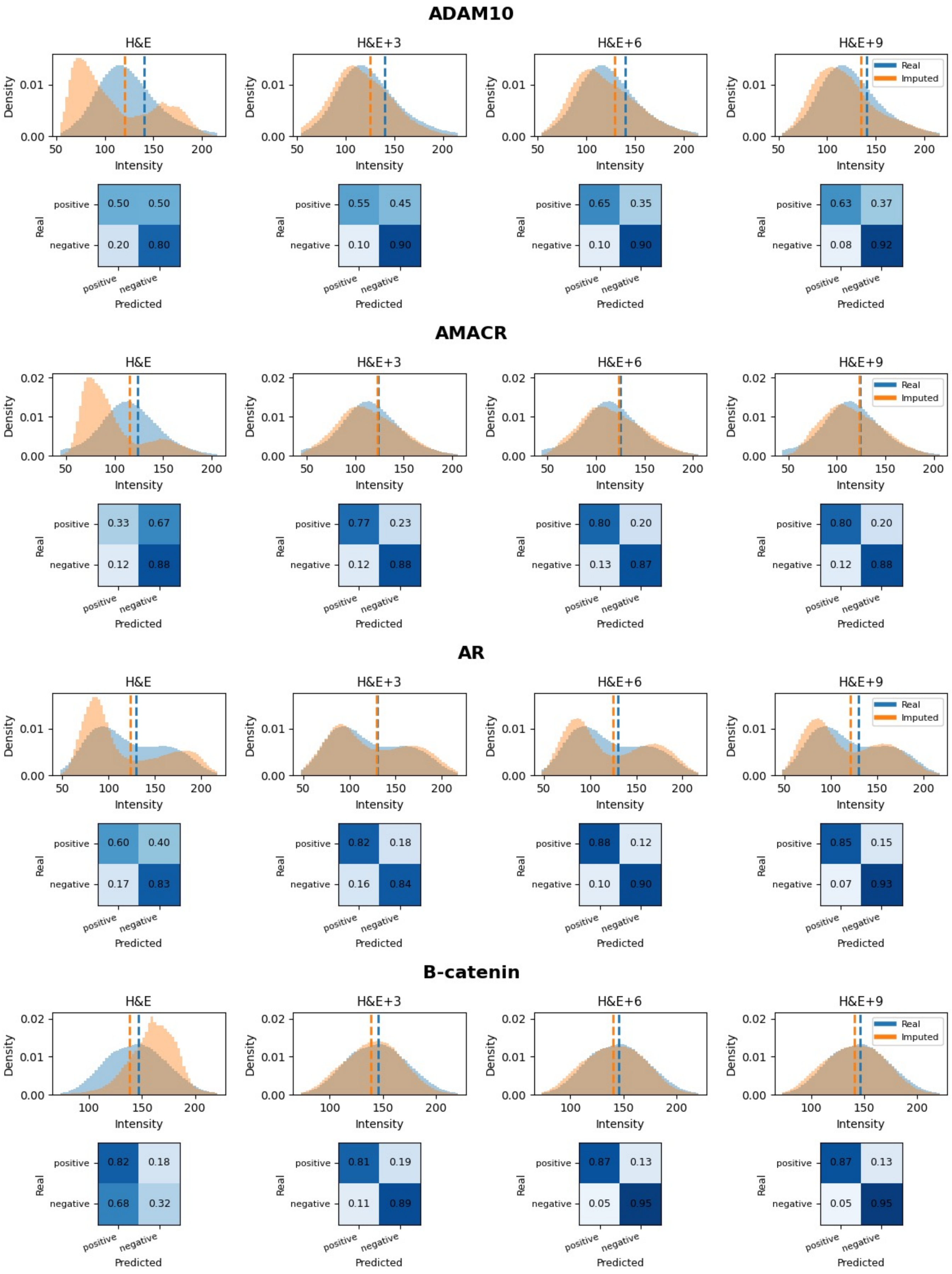

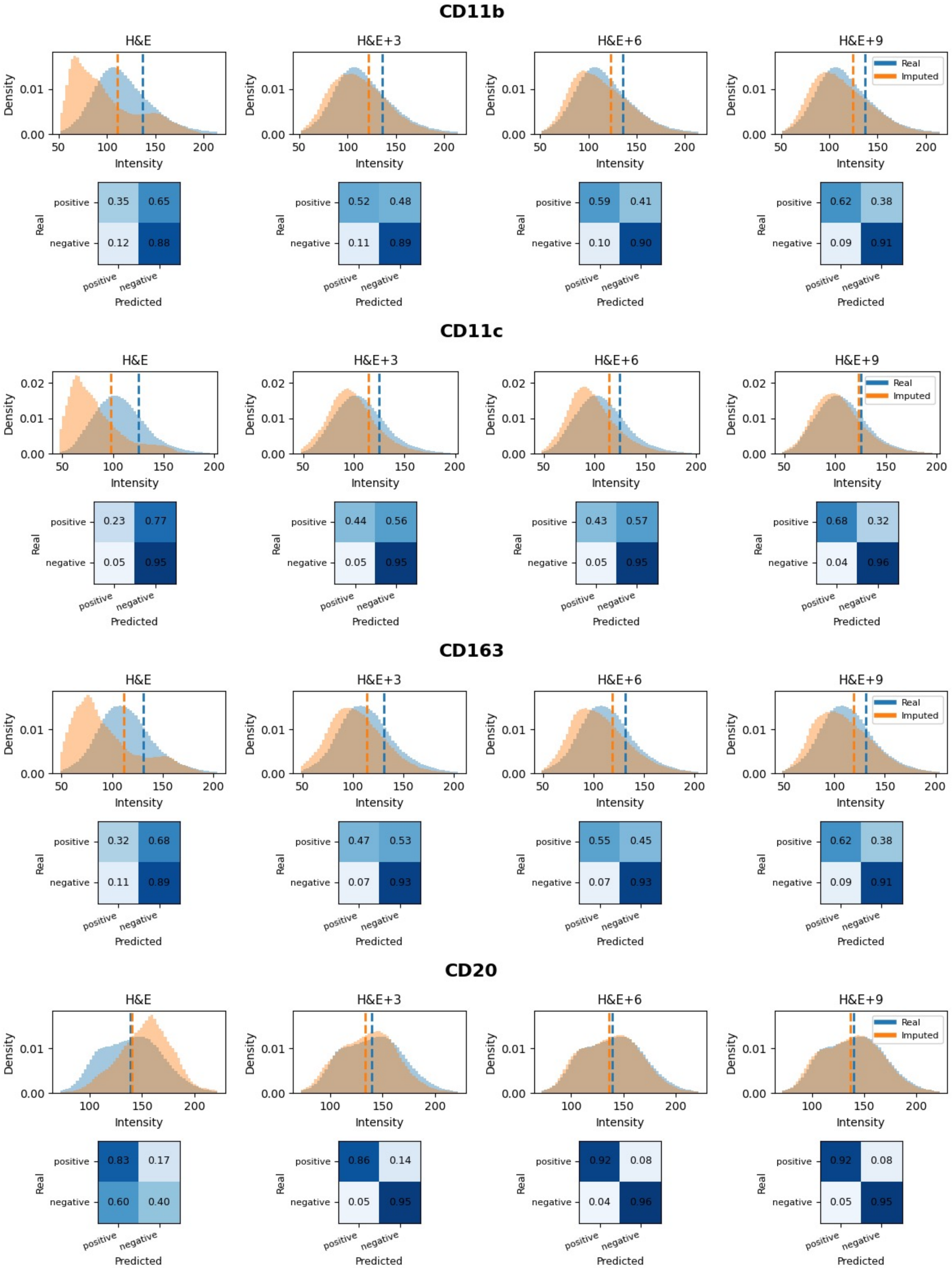
CD11b
H&E
H&E+3
H&E+6
H&E+9
Real
Imputed
Density
Intensity
positive
negative
Real
Predicted
0.35 0.65
0.12 0.88
0.52 0.48
0.11 0.89
0.59 0.41
0.10 0.90
0.62 0.38
0.09 0.91
CD11c
0.23 0.77
0.05 0.95
0.44 0.56
0.05 0.95
0.43 0.57
0.05 0.95
0.68 0.32
0.04 0.96
CD163
0.32 0.68
0.11 0.89
0.47 0.53
0.07 0.93
0.55 0.45
0.07 0.93
0.62 0.38
0.09 0.91
CD20
0.83 0.17
0.60 0.40
0.86 0.14
0.05 0.95
0.92 0.08
0.04 0.96
0.92 0.08
0.05 0.95

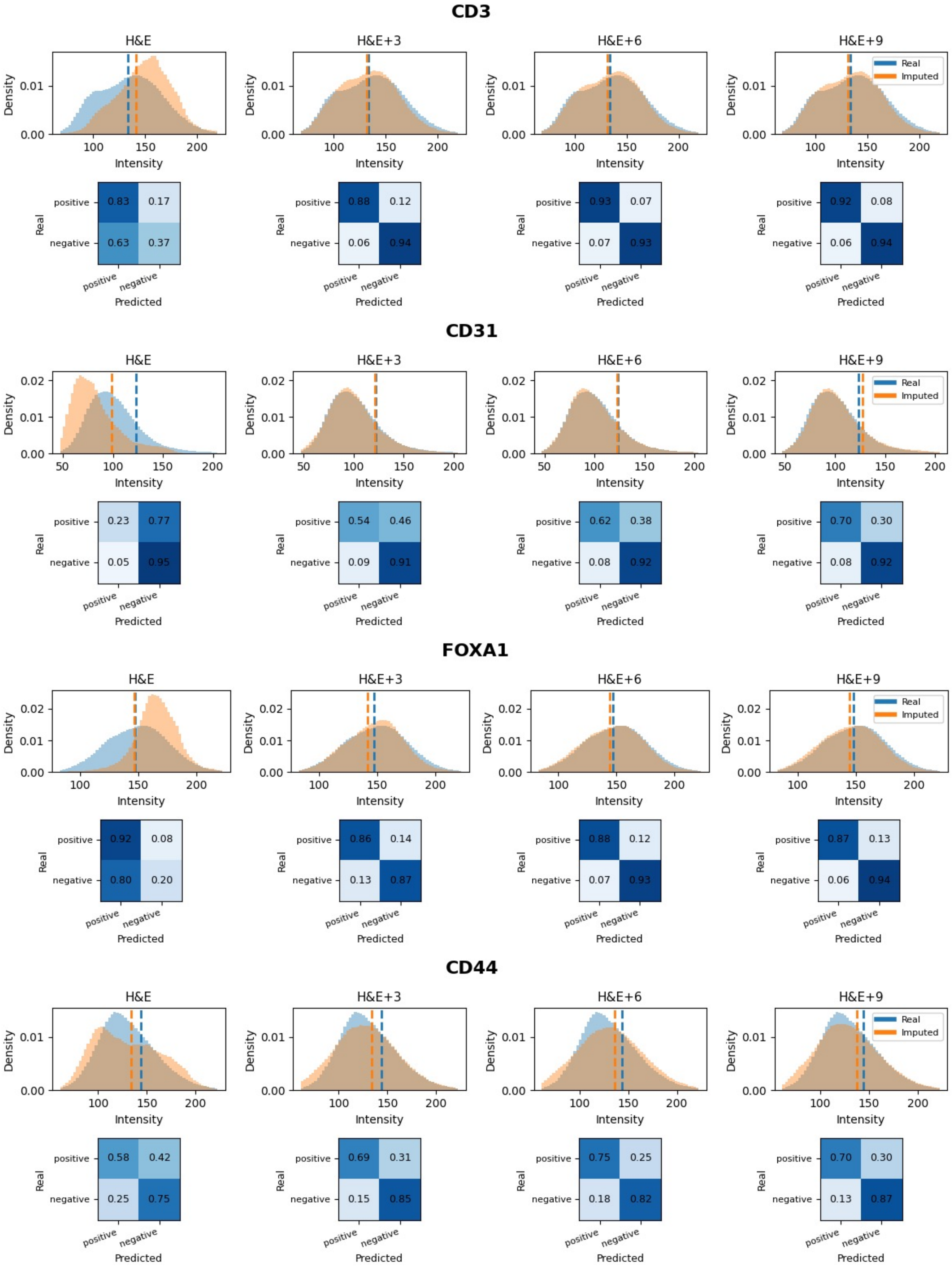

CD3
H&E
H&E+3
H&E+6
H&E+9
Density
Intensity
Real
Imputed
positive 0.83 0.17
negative 0.63 0.37
positive 0.88 0.12
negative 0.06 0.94
positive 0.93 0.07
negative 0.07 0.93
positive 0.92 0.08
negative 0.06 0.94
Real
Predicted
CD31
positive 0.23 0.77
negative 0.05 0.95
positive 0.54 0.46
negative 0.09 0.91
positive 0.62 0.38
negative 0.08 0.92
positive 0.70 0.30
negative 0.08 0.92
FOXA1
positive 0.92 0.08
negative 0.80 0.20
positive 0.86 0.14
negative 0.13 0.87
positive 0.88 0.12
negative 0.07 0.93
positive 0.87 0.13
negative 0.06 0.94
CD44
positive 0.58 0.42
negative 0.25 0.75
positive 0.69 0.31
negative 0.15 0.85
positive 0.75 0.25
negative 0.18 0.82
positive 0.70 0.30
negative 0.13 0.87

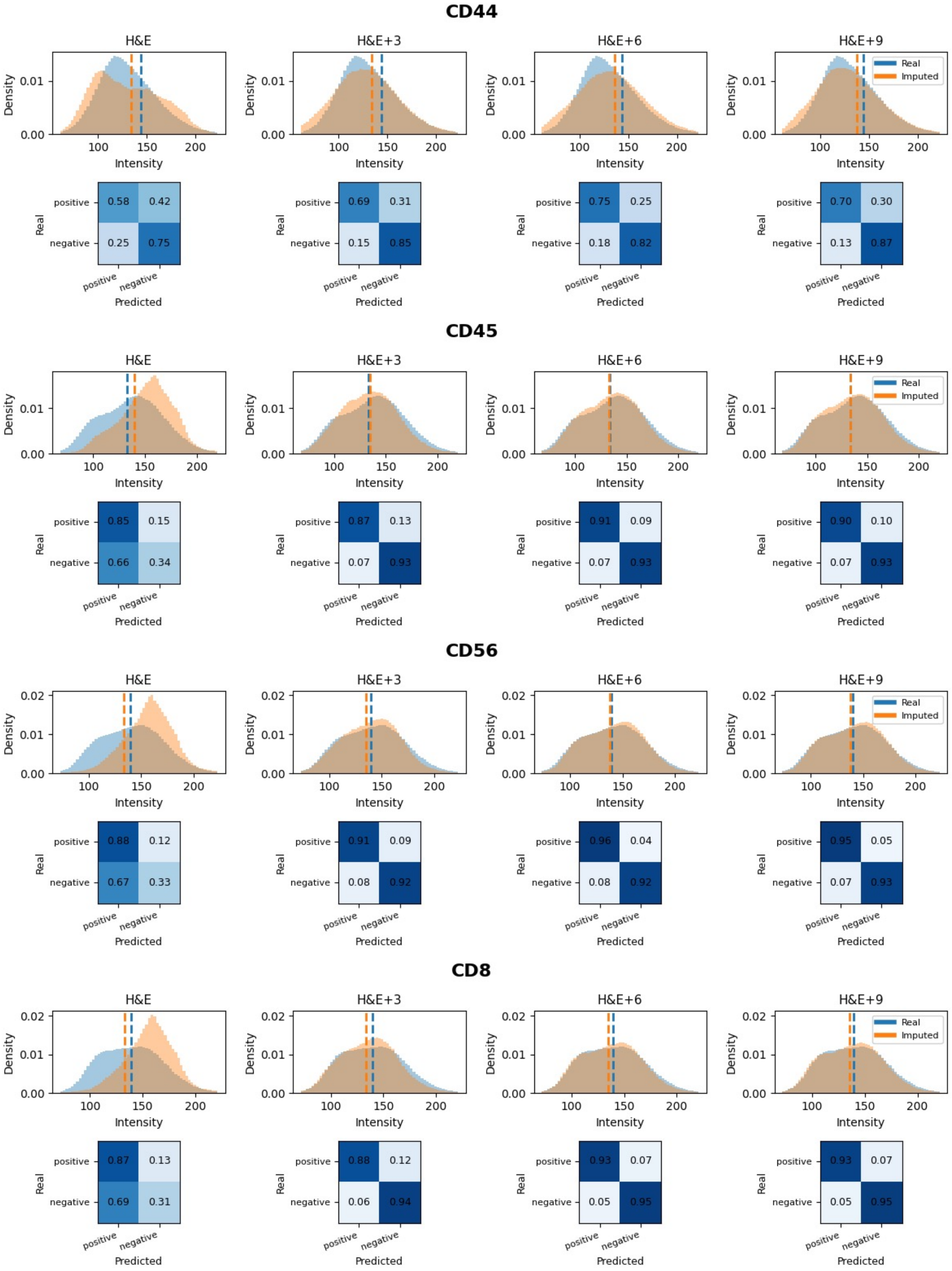
CD44
H&E
H&E+3
H&E+6
H&E+9
Density
Intensity
Real
Imputed
positive
negative
Predicted
0.58 0.42
0.25 0.75
0.69 0.31
0.15 0.85
0.75 0.25
0.18 0.82
0.70 0.30
0.13 0.87
CD45
0.85 0.15
0.66 0.34
0.87 0.13
0.07 0.93
0.91 0.09
0.07 0.93
0.90 0.10
0.07 0.93
CD56
0.88 0.12
0.67 0.33
0.91 0.09
0.08 0.92
0.96 0.04
0.08 0.92
0.95 0.05
0.07 0.93
CD8
0.87 0.13
0.69 0.31
0.88 0.12
0.06 0.94
0.93 0.07
0.05 0.95
0.93 0.07
0.05 0.95

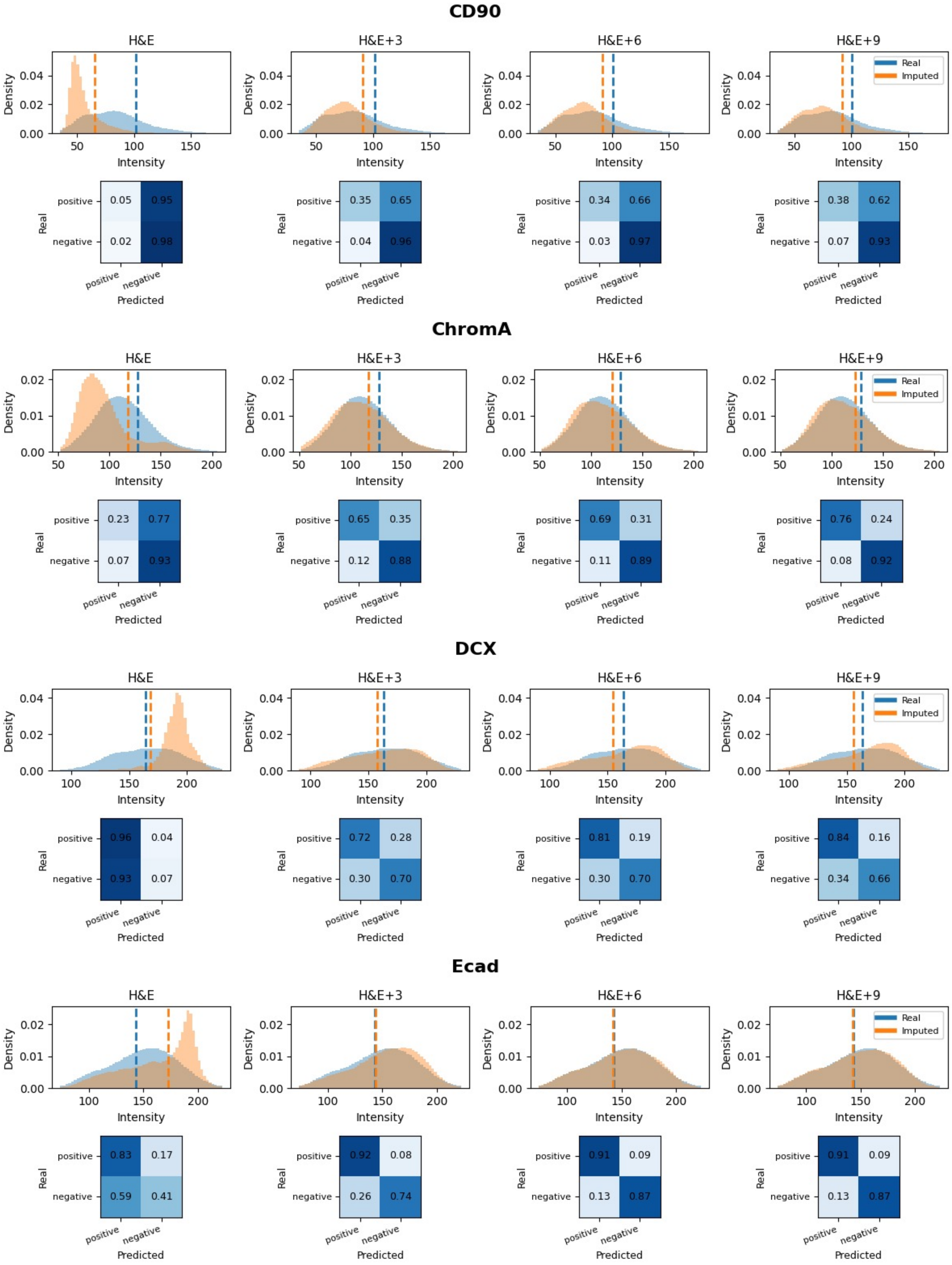
CD90
H&E
H&E+3
H&E+6
H&E+9
Real
Imputed
Density
Intensity
positive
negative
Real
Predicted
0.05 0.95
0.02 0.98
0.35 0.65
0.04 0.96
0.34 0.66
0.03 0.97
0.38 0.62
0.07 0.93
ChromA
0.23 0.77
0.07 0.93
0.65 0.35
0.12 0.88
0.69 0.31
0.11 0.89
0.76 0.24
0.08 0.92
DCX
0.96 0.04
0.93 0.07
0.72 0.28
0.30 0.70
0.81 0.19
0.30 0.70
0.84 0.16
0.34 0.66
Ecad
0.83 0.17
0.59 0.41
0.92 0.08
0.26 0.74
0.91 0.09
0.13 0.87
0.91 0.09
0.13 0.87

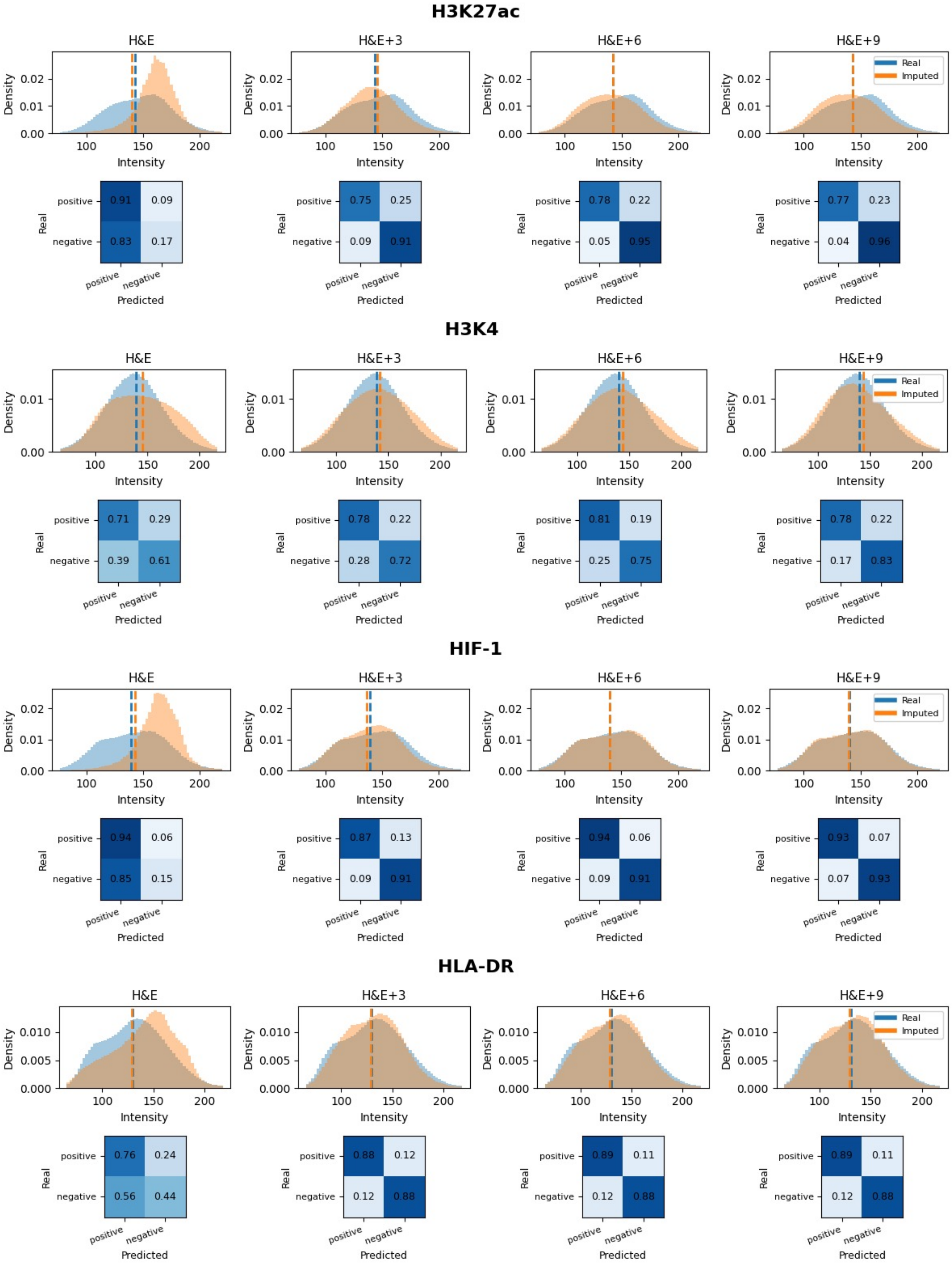
H3K27ac
H&E
H&E+3
H&E+6
H&E+9
Density
Intensity
Real
Imputed
positive
negative
Real
Predicted
0.91 0.09
0.83 0.17
0.75 0.25
0.09 0.91
0.78 0.22
0.05 0.95
0.77 0.23
0.04 0.96
H3K4
H&E
H&E+3
H&E+6
H&E+9
0.71 0.29
0.39 0.61
0.78 0.22
0.28 0.72
0.81 0.19
0.25 0.75
0.78 0.22
0.17 0.83
HIF-1
H&E
H&E+3
H&E+6
H&E+9
0.94 0.06
0.85 0.15
0.87 0.13
0.09 0.91
0.94 0.06
0.09 0.91
0.93 0.07
0.07 0.93
HLA-DR
H&E
H&E+3
H&E+6
H&E+9
0.76 0.24
0.56 0.44
0.88 0.12
0.12 0.88
0.89 0.11
0.12 0.88
0.89 0.11
0.12 0.88

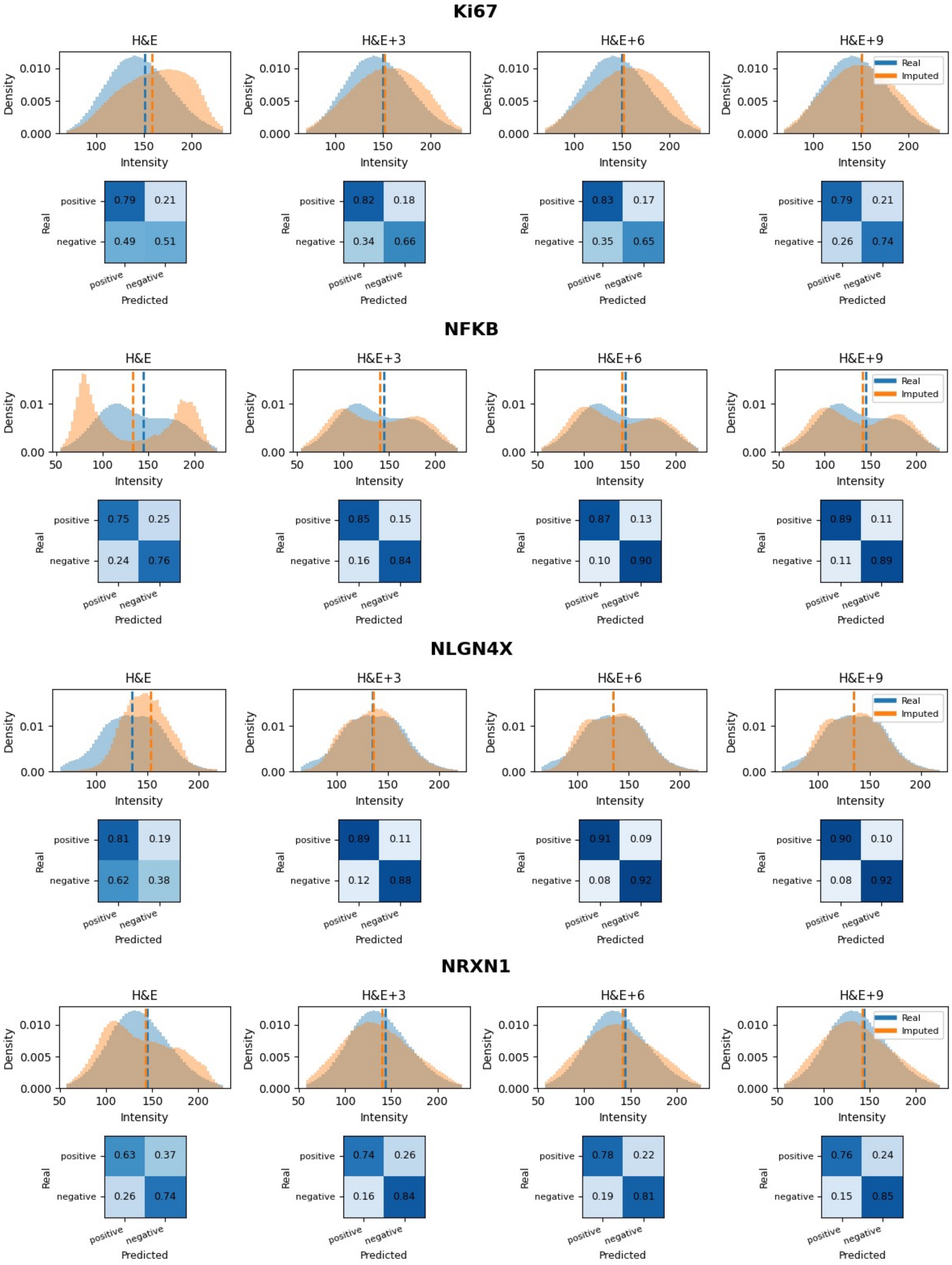
Ki67
H&E
H&E+3
H&E+6
H&E+9
Real
Imputed
Density
Intensity
Real
Predicted
positive
negative
0.79 0.21
0.49 0.51
0.82 0.18
0.34 0.66
0.83 0.17
0.35 0.65
0.79 0.21
0.26 0.74
NFKB
0.75 0.25
0.24 0.76
0.85 0.15
0.16 0.84
0.87 0.13
0.10 0.90
0.89 0.11
0.11 0.89
NLGN4X
0.81 0.19
0.62 0.38
0.89 0.11
0.12 0.88
0.91 0.09
0.08 0.92
0.90 0.10
0.08 0.92
NRXN1
0.63 0.37
0.26 0.74
0.74 0.26
0.16 0.84
0.78 0.22
0.19 0.81
0.76 0.24
0.15 0.85

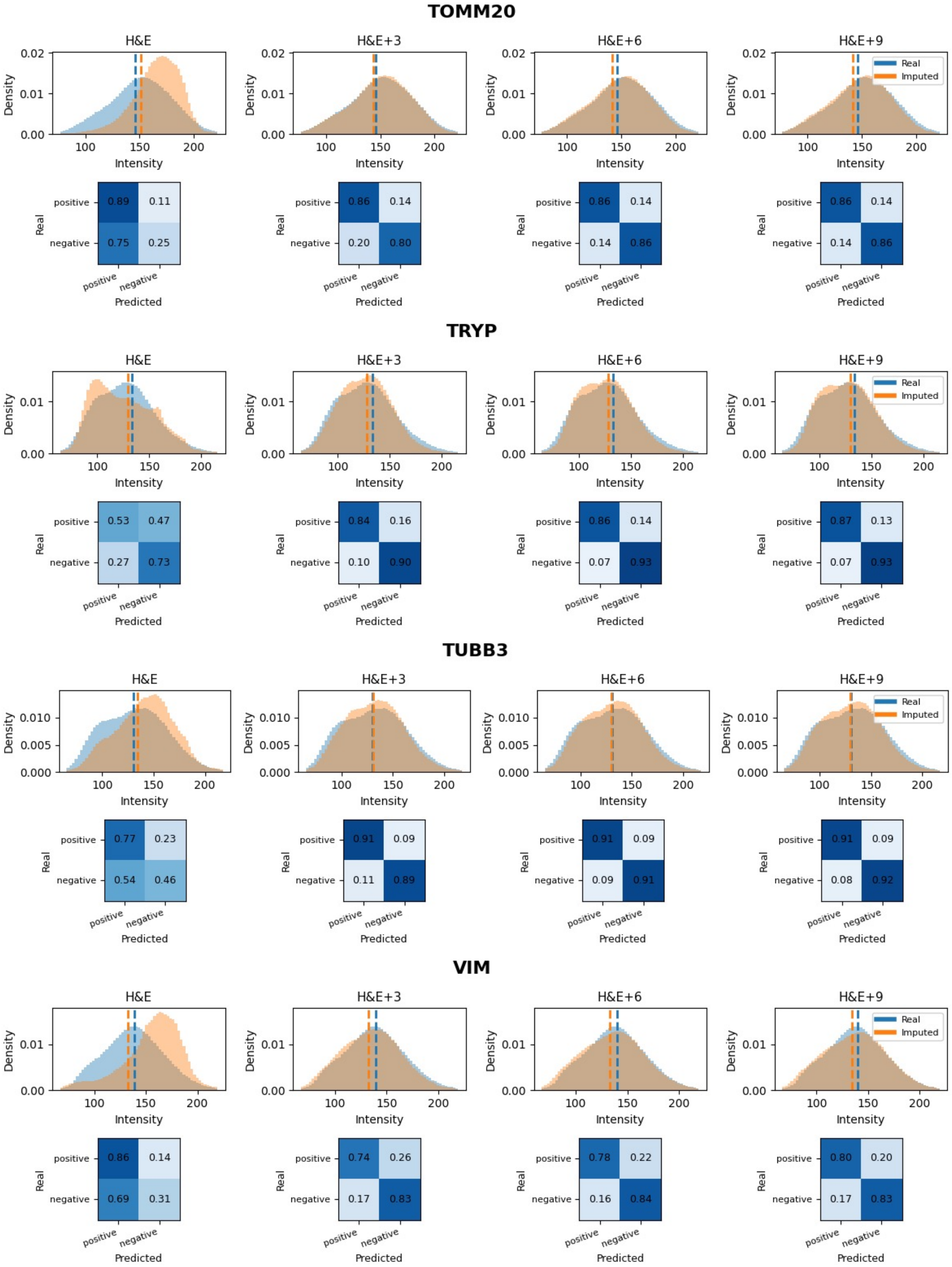
TOMM20
H&E
H&E+3
H&E+6
H&E+9
Density
Intensity
Real
Imputed
positive
negative
Predicted
0.89
0.11
0.75
0.25
0.86
0.14
0.20
0.80
0.86
0.14
0.14
0.86
0.86
0.14
0.14
0.86
TRYP
0.53
0.47
0.27
0.73
0.84
0.16
0.10
0.90
0.86
0.14
0.07
0.93
0.87
0.13
0.07
0.93
TUBB3
0.77
0.23
0.54
0.46
0.91
0.09
0.11
0.89
0.91
0.09
0.09
0.91
0.91
0.09
0.08
0.92
VIM
0.86
0.14
0.69
0.31
0.74
0.26
0.17
0.83
0.78
0.22
0.16
0.84
0.80
0.20
0.17
0.83

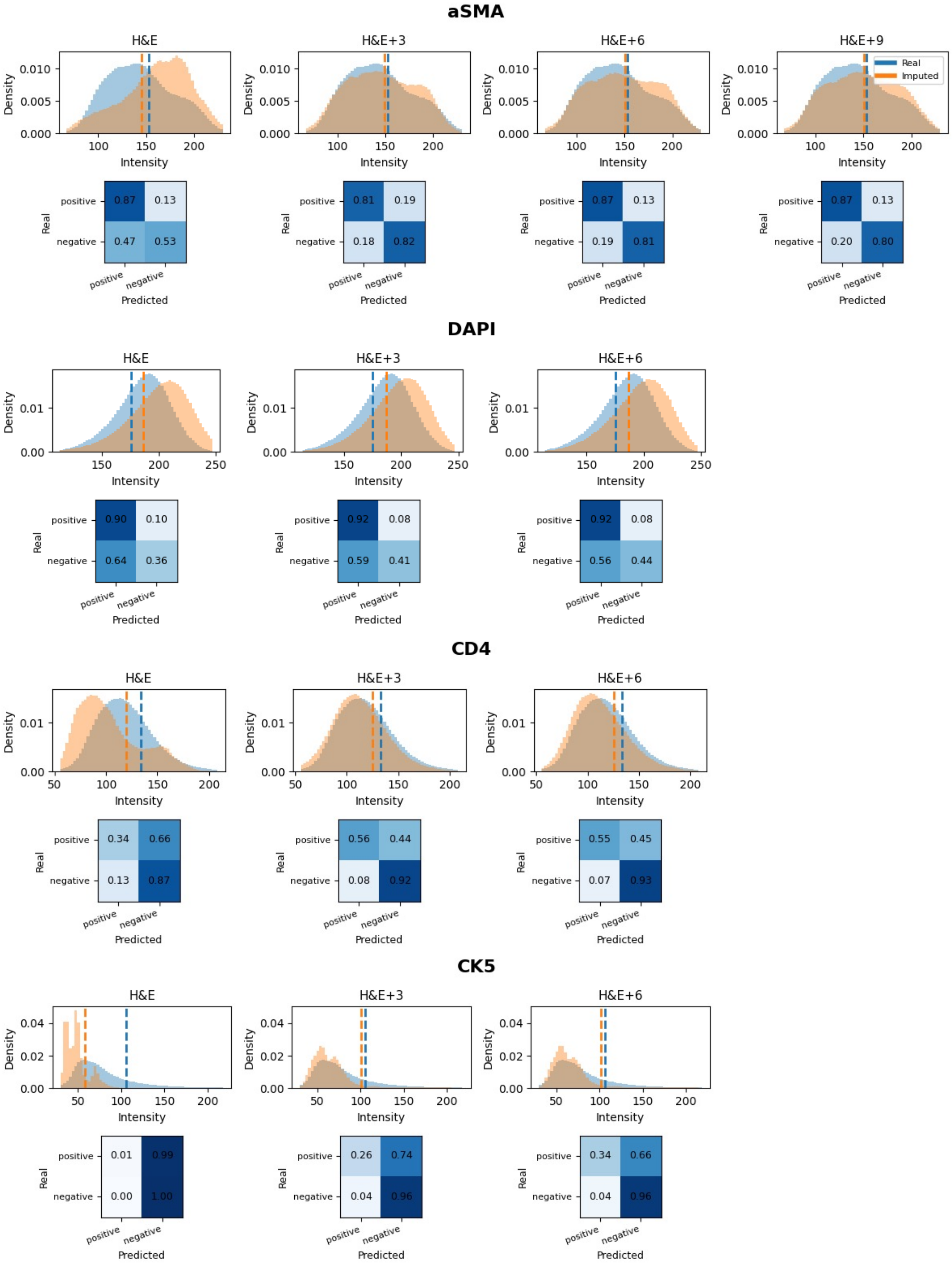
aSMA
H&E
H&E+3
H&E+6
H&E+9
Real
Imputed
Density
Intensity
Real
Predicted
positive
negative
0.87
0.13
0.47
0.53
0.81
0.19
0.18
0.82
0.87
0.13
0.19
0.81
0.87
0.13
0.20
0.80
DAPI
0.90
0.10
0.64
0.36
0.92
0.08
0.59
0.41
0.92
0.08
0.56
0.44
CD4
0.34
0.66
0.13
0.87
0.56
0.44
0.08
0.92
0.55
0.45
0.07
0.93
CK5
0.01
0.99
0.00
1.00
0.26
0.74
0.04
0.96
0.34
0.66
0.04
0.96

#### GZMB

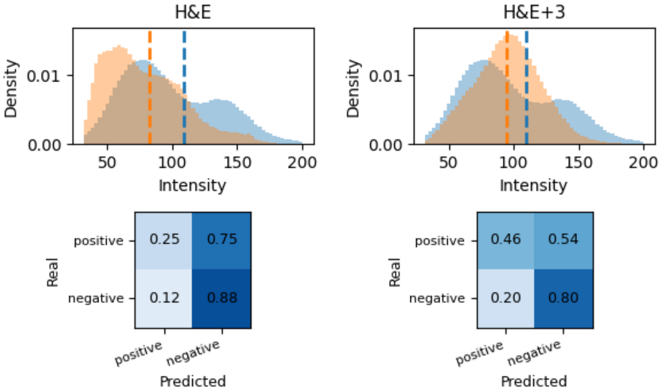


## CK8

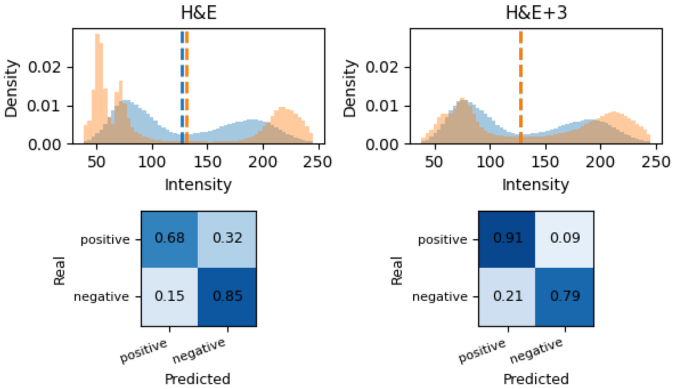


#### NLGN1

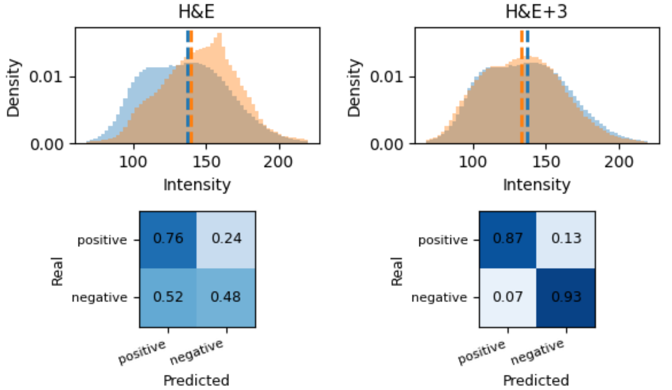


## CD68

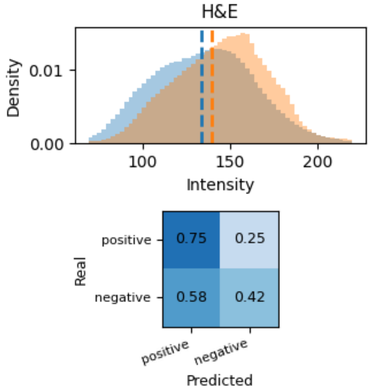

## TH

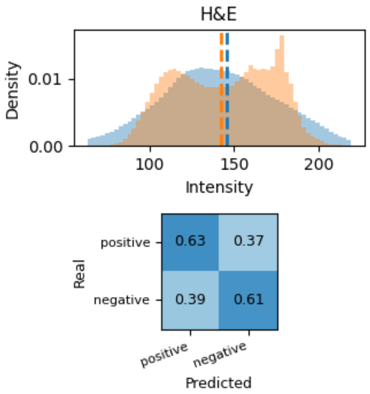
H&E
Density
0.01
0.00
100
150
200
Intensity
positive
negative
Real
0.63
0.37
0.39
0.61
positive
negative
Predicted

#### ERG

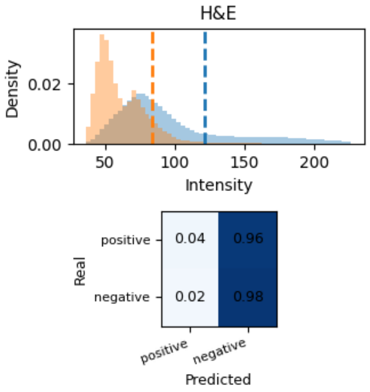
H&E
Density
0.02
0.00
50
100
150
200
Intensity
positive
negative
Real
0.04
0.96
0.02
0.98
positive
negative
Predicted

### Supplementary Figure 5

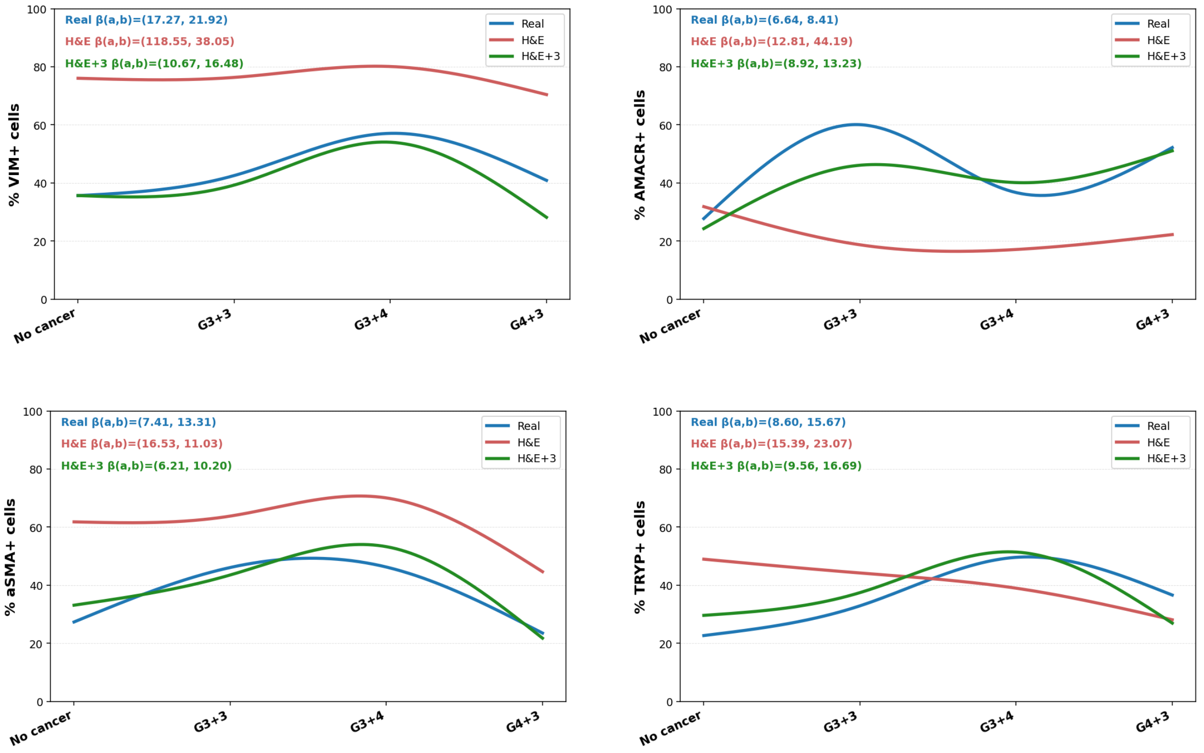

### Supplementary Figure 6

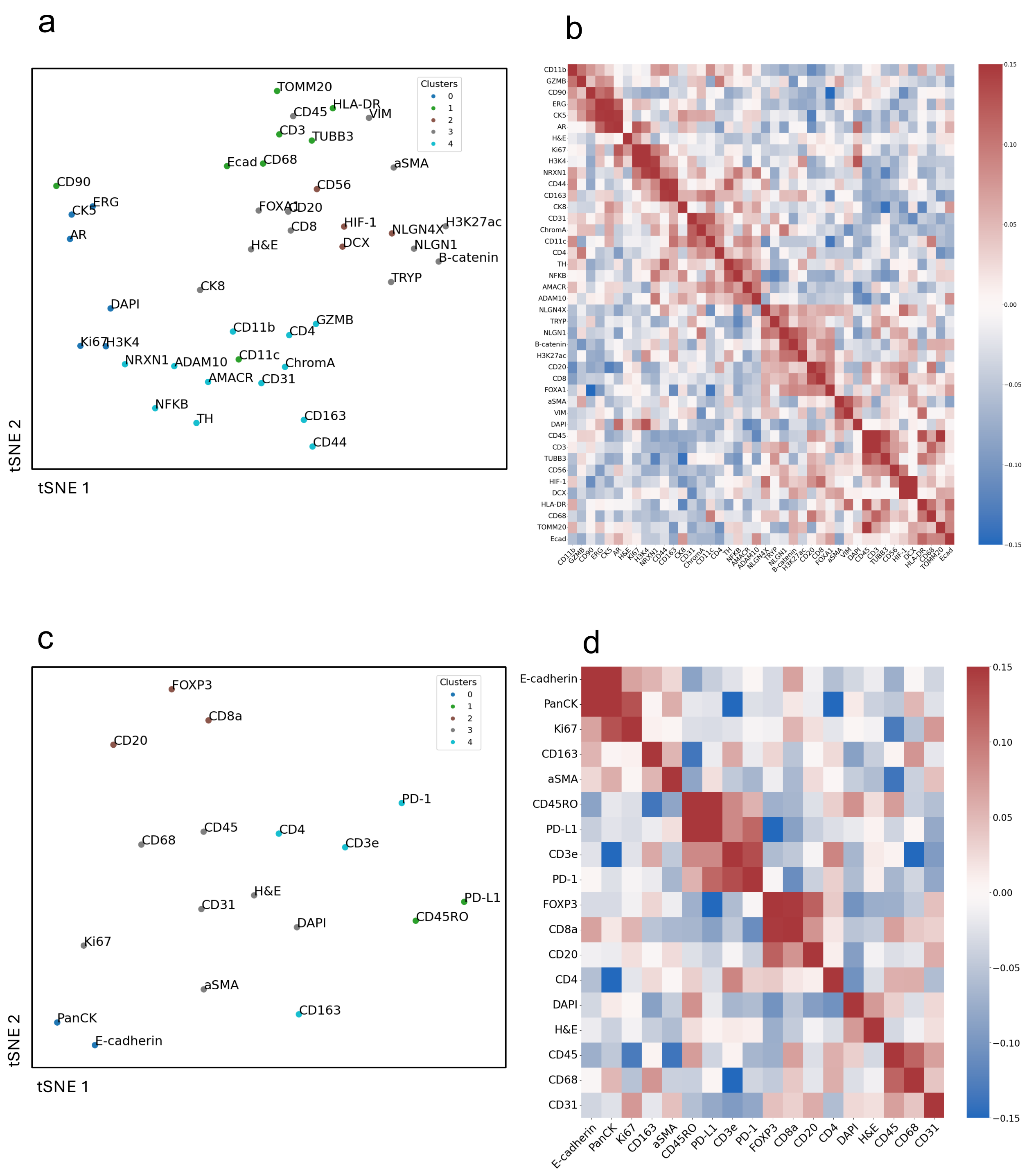

### Supplementary Figure 7

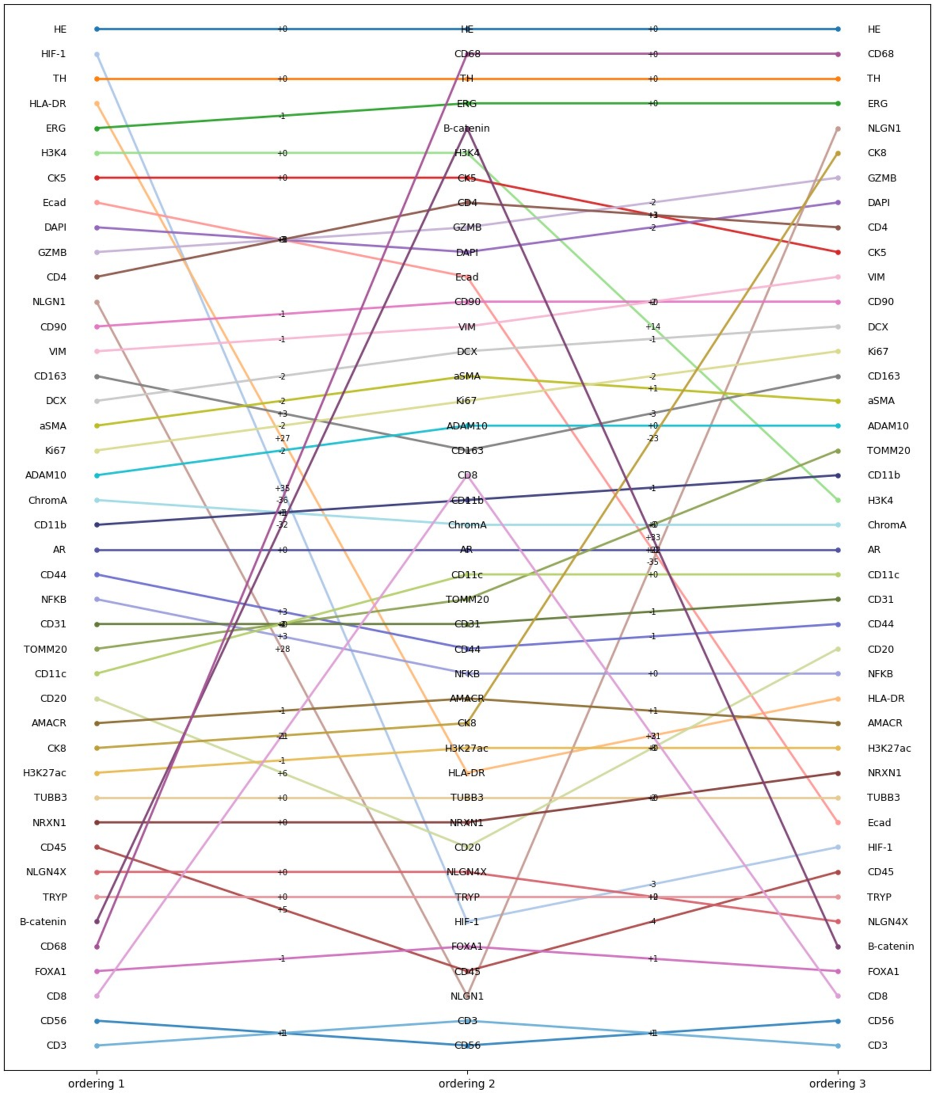

### Supplementary Table 1

| Model | Spearman | SSIM |
|---|---|---|
| IF-256 | 0.98 | 0.77 |
| IF-1024 | 0.95 | 0.82 |
| IF-HE-1024 | 0.98 | 0.8 |
| IF-HE-256 | 0.98 | 0.78 |
| IF-HE-256,256 | 0.96 | 0.87 |
| IF-HE-128,256 | 0.94 | 0.84 |
| IF-HE-128,128 | 0.52 | 0.78 |

| CHANNEL WISE SPEARMAN | | | | | | | | | | | | | | | | | | | | |
|---|---|---|---|---|---|---|---|---|---|---|---|---|---|---|---|---|---|---|---|---|
| Model | DAPI | CD31 | CD45 | CD68 | CD4 | FOXP3 | CD8A | CD45RO | CD20 | PD-L1 | CD3e | CD163 | E-Cadherin | PD-1 | Ki67 | PanCK | aSMA | E | H | Average |
| **IF HE 256,256** | 0.8949 | 0.9947 | 0.9220 | 0.9715 | 0.9929 | 0.9813 | 0.9863 | 0.9963 | 0.9937 | 0.9970 | 0.9932 | 0.9895 | 0.9015 | 0.9941 | 0.9896 | 0.7037 | 0.9714 | 0.9714 | 0.9772 | 0.97 |
| **IF HE 128, 256** | 0.9609 | 0.9884 | 0.9835 | 0.9797 | 0.9859 | 0.9770 | 0.9757 | 0.9910 | 0.9838 | 0.9903 | 0.9876 | 0.9891 | 0.7695 | 0.9895 | 0.9862 | 0.6945 | 0.9574 | 0.9553 | 0.8050 | 0.94 |
| **IF HE 128,128** | 0.0388 | 0.0588 | 0.6588 | 0.5388 | 0.0892 | 0.492 | 0.9892 | 0.715 | 0.1584 | 0.9888 | 0.5693 | 0.8222 | 0.6470 | 0.5126 | 0.9823 | 0.1584 | 0.9892 | 0.1745 | 0.2735 | 0.52 |
